# Climate warming and urbanization may expand dengue transmission risk in California

**DOI:** 10.1101/2025.11.14.688544

**Authors:** Lisa I. Couper, Terrell J Sipin, Samantha Sambado, Zoe Rennie, Kyle M. Shanebeck, Kelsey P. Lyberger, Philip P.A. Collender, Van Ngo, Justin V. Remais, Andrew J. MacDonald

## Abstract

**Background:** While primarily a disease of tropical and subtropical regions, dengue outbreaks are increasing in non-endemic regions due to environmental change and increasing travel and trade. For these non-endemic regions, estimating the risk of dengue is challenging as transmission is driven by both local environmental conditions and the introduction of viremic travelers. In this study, we aimed to estimate current and future dengue risk in California, USA—a region that has recently experienced its first cases of locally-acquired dengue.

**Methods:** We modeled dengue risk as the product of three key components needed for local transmission—vector presence, temperature-suitability for pathogen transmission, and viral introductions via travel-associated cases—estimated using vector and case surveillance, sociodemographic, and environmental data. We estimated risk for locations and months where local transmission was reported in 2023-2024 to define a ‘threshold’ level of risk. We then projected monthly, census tract-level risk under both current conditions and future scenarios of climate warming and urban expansion.

**Findings:** Approximately 18.2 million (95% CI: 17.9-18.3) California residents—primarily in the Central Valley and the Los Angeles and San Diego metropolitan areas—currently live in areas where peak monthly dengue risk exceeds levels estimated during observed local transmission. Under moderate scenarios of climate warming and urban expansion, an additional 4.1 million (95% CI: 3.7-4.6) California residents may be at risk by mid-century, with the largest increase in risk estimated for September and for the Sacramento Valley and coastal southern California regions. Outside the summer months and beyond the Central Valley and southern California, current and future risk remains low due to one or more major bottlenecks to transmission.

**Interpretation:** Our study identifies the specific regions and months conducive to dengue transmission in the non-endemic setting of California. At present, this covers a substantial portion of the state and is projected to expand under on-going climate warming and urbanization. Our results underscore the need for sustained vector control, and timely detection and management of travel-associated cases.

**Research in Context:** *Evidence before this study:* Dengue is considered endemic in over 125 countries and rapidly expanding its range, aided by climate warming, urbanization, and global travel and trade. Estimating transmission risk in newly emerging regions is critical for public health preparedness and depends on both local environmental conditions and the introduction of viremic travelers. We searched PubMed from database inception to May 8, 2025, for articles published in English using search terms “dengue”, “model”, “non-endemic”, and their common textual variants. We identified 75 relevant studies modeling dengue transmission risk in non-endemic settings. However, nearly all were focused on one or two major determinants of transmission (eg, climate, vector population dynamics, or case importations) and/or did not include future projections. We found no studies that developed and validated a model of dengue transmission risk in non-endemic settings that incorporated vector, pathogen, and human suitability factors, and applied this model to project future risk.

*Added value of this study:* This study provides a novel approach to model dengue transmission risk in emerging regions that integrates the major factors driving transmission—vector presence, temperature suitability, and travel-associated cases. We apply this model to California—an emerging center of transmission risk in the continental USA—to identify the times and regions where risk exceeds levels observed during recent local transmission. We found that approximately 18.2 million California residents may be at risk based on this threshold, with an additional 4.1 million potentially at risk by mid-century under a moderate scenario of warming and urban expansion.

*Implications of all the available evidence:* Our study identifies the hotspots of dengue transmission risk at a fine spatial and temporal resolution (census tract, month) in a highly populous and globally-connected region of emerging dengue risk. These risk estimates, and the regionally-specific bottlenecks to transmission that we identify can inform targeted disease surveillance and prevention strategies. Further, our findings have implications for other emerging regions including the southern USA and southern Europe, suggesting that the risk of local dengue transmission may increase under ongoing climate warming, urbanization, and global travel.

## Introduction

Ongoing and increasingly rapid environmental change and globalization are having profound effects on both ecological systems and human health (1–3). In particular, these anthropogenic changes are contributing to the rise in emerging infectious diseases, especially at higher latitudes where climate warming is facilitating the expansion of arthropod vectors of disease (4–6). For example, aided by climate change and globalization, the global burden of dengue has increased tenfold in recent decades—from half a million reported cases in 2000 to over 10 million in 2023—posing a major threat to public health (7–10). While over 125 countries are now considered endemic, climate warming is predicted to substantially expand dengue transmission suitability in currently non-endemic regions including in Europe, Canada, Argentina, southern Africa, and the United States (10–12). In the continental USA, outbreaks of dengue have recently occurred in southern states including Florida, Texas, Arizona, and California (13–15). As outbreaks in these regions are expected to become more frequent (16), identifying the areas of greatest risk for ongoing transmission and preparing targeted interventions is critical for protecting public health.

As dengue is transmitted by ectothermic mosquito vectors, its emergence in non-endemic regions is facilitated by warming temperatures increasing the suitability for disease transmission (17–21). In particular, at cold temperatures, transmission is inhibited due to low performance of mosquito and/or pathogen life history traits including survival, development, and reproduction (22–26). As temperatures rise above these lower limits for trait performance, transmission becomes possible, and small increases in temperature can have large impacts on mosquito-borne disease transmission and incidence (27,28). Notably, dengue has one of the highest optimal temperatures for transmission of all mosquito-borne diseases, with peak transmission by *Aedes aegypti* occurring around 29°C (28.4-29.8°C estimated 95% CI) (25,29). This makes dengue a focal disease of concern under climate warming, as rising temperatures may push more regions into this optimal transmission window.

In addition to increasing climate suitability, international trade and travel have facilitated the spread of dengue in non-endemic regions (30–32). International trade, particularly the shipping of used tires, has been linked to the introduction of *Aedes* vectors to several countries including the USA, Italy, Albania, and South Africa (33–41). Once introduced, *Ae. aegypti* and *Ae. albopictus*—the two main vectors of dengue virus (DENV) (42)—readily spread and become established owing to their ecological plasticity (8,18,43–47). While introduced *Aedes* populations are unlikely to be infected with DENV, infected travelers returning from endemic countries are a well-known, and potentially increasing, source of DENV spread to non-endemic regions (31,48–50). Indeed, DENV infections are common among international travelers, and epidemics in hyper-endemic regions are often reflected in travel-associated case burdens in non-endemic regions (30,47,51–54).

Here we develop a novel approach to estimate local dengue risk in emerging regions by combining information on the major factors needed for transmission: vector presence, suitable temperatures for pathogen transmission processes, and viral introductions via travel-associated cases. We apply this approach to estimate transmission risk in California, USA—an emerging center of dengue risk with a robust, highly coordinated mosquito-borne disease surveillance program. California reported its first two cases of locally-acquired dengue in Los Angeles County in 2023, and several small outbreaks across southern California in 2024. Both *Ae. aegypti* and *Ae. albopictus* were first identified in the state in the early 2010s, and *Ae. aegypti* in particular has since rapidly expanded, establishing distinct breeding populations in northern, central, and southern California (14,55–57). Further, given its large population size (nearly 40 million residents), introductions of viremic hosts occur relatively frequently in California, with approximately 140 cases of travel-associated dengue reported each year—likely an underestimate of the true case count (14,58). We estimated monthly dengue risk across the state under current and future projections of climate warming and urban expansion to identify the locations and times where heightened preventative measures may be most needed.

## Methods

We estimated the risk of local dengue transmission for a given month and location by estimating the following factors necessary for transmission: 1) the presence of competent vectors (here expressed as a probability); 2) suitable temperatures for pathogen transmission (expressed as a scaled version of *R*_0_(T)); and 3) viral introductions (expressed as the number of travel-associated dengue cases per year) (Figure 1).

**Figure 1.**
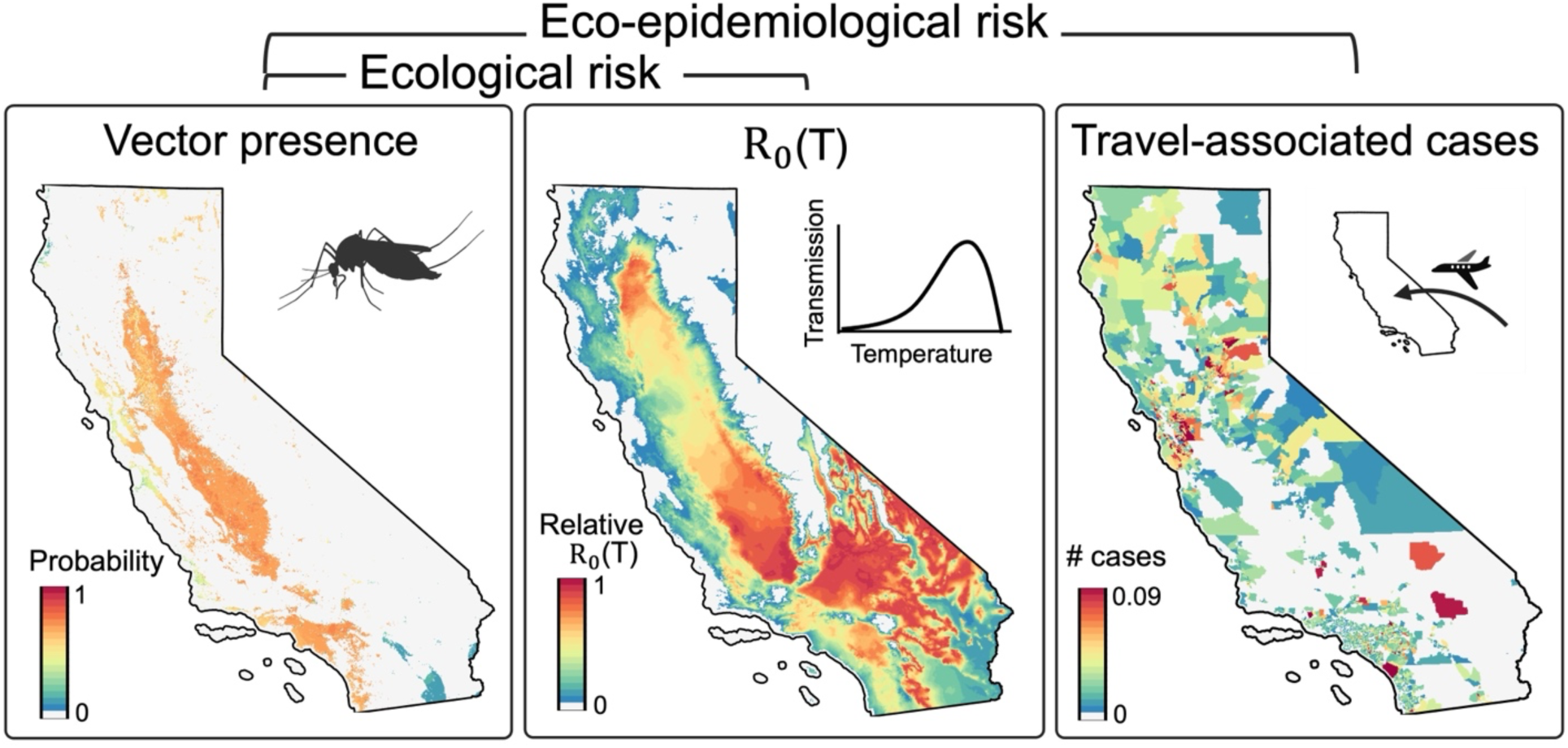
Factors included in local dengue transmission risk indices. The first risk index, ‘ecological risk’, is estimated based on the probability of *Aedes aegypti* presence and the temperature-based suitability for dengue transmission (ie, *R*_0_(T) scaled from 0 to 1). The second index, ‘eco-epidemiological risk’, additionally incorporates an estimate of the number of travel-associated cases for a given month and census tract (here, ranging from 0 to 0.09—the lowest and highest values of travel-associated cases estimated across all tract-months, respectively). Estimates in the example above are for August 2023.

### Vector species distribution model

To estimate the probability of competent vector presence across California, we developed monthly species distribution models (SDMs) using two common modeling approaches—Maximum Entropy (MaxEnt) and Extreme Gradient Boosting (XGBoost) (59). Occurrence data were drawn from the California Vectorborne Disease Surveillance System (CalSurv) trap surveillance data (through Data Use Request 000050). These data included trap location (latitude and longitude), collection date, trap type, and the number of nights deployed. While this *Ae. aegypti* surveillance data are available from 2013 (the year of first detection in the state) to 2023, we constrained our model to later years (2016–2023) to avoid attributing vector absences to dispersal limitation during early invasion. Although *Ae. albopictus* is also a competent vector for dengue and has become established in California (57,60,61), we focus on *Ae. aegypti* due to its broader distribution, more frequent detection, and generally higher DENV competence (62–64).

Both SDM modeling approaches rely on comparing values of environmental predictors between species occurrences and background points (or ‘pseudo-absences’) that represent the landscape across which the species could be present. Here, we generated background points for *Ae. aegypti* using a two-step approach. First, we used monthly occurrences of adult *Culex quinquefasciatus* from the CalSurv trap surveillance data, as an indicator of potentially suitable locations where mosquito surveillance is occurring, but *Ae. aegypti* is not identified. Since vector surveillance is spatially biased and not randomly assigned across the state, we then used a temperature suitability mask to randomly sample pseudo-absence points, weighted toward regions known to be unsuitable based on physiological constraints to *Ae. aegypti* population persistence (following methods in Kraemer et al. 2015) (45) (Supplementary Methods). For both modeling approaches, the climate variables considered as predictors included mean temperature, total precipitation, and terrestrial water storage—the total amount of water on and below the land surface. Monthly values of all climate variables were obtained from the California Basin Characterization Model (CA BCM) given its high spatial resolution (270 m) (65). Because invasive *Ae. aegypti* populations are adapted to urban settings, with larvae developing in artificial containers, and adults feeding primarily on humans, we masked outputs from both modeling approaches based on land cover types that are highly unlikely to harbor *Ae. aegypti* rather than including land cover in models directly (45,66) (Supplementary Methods).

Finally, to ensure our results were robust to modeling approaches and assumptions, we combined results from the two SDMs (ie, the MaxEnt and XGBoost models) into a meta-learner (67–69). The meta-learner is trained on predictions of the individual learners, thereby learning from the mistakes of each base learner and adopting their strengths. Our meta-learner used a random forest algorithm tuned through spatiotemporal cross-validation and achieved slightly better performance than either of the individual approaches alone (Supplementary Methods).

To estimate future vector presence probabilities, we obtained climate projections from the Model for Interdisciplinary Research on Climate (MIROC) under Representative Concentration Pathways (RCPs) 4.5, 6.0, and 8.5 for the period from 2010-2020 (‘current’), 2040-2050 (‘mid-century’), and 2090-2100 (‘end-of-century’). Data on current and future land cover were obtained from the USGS Forecasting Scenarios of land use (FORE-SCE) model under the A2 scenario projection and available at a 250 m resolution (70) (Supplementary Methods, Supplementary Figure S1).

### Temperature-based transmission suitability

As temperature imposes physiological limits on both mosquito and viral trait performance, we estimated the temperature-based transmission suitability of dengue following methods from Mordecai et al. 2017 (25). Therein, data from laboratory experiments on vector and viral trait performance at different constant temperatures were used to estimate continuous thermal response curves. Thermal responses for all temperature-dependent traits involved in transmission, including vector survival, development rates, biting rate, adult lifespan, fecundity, competence, and viral extrinsic incubation period, were then integrated into an estimate of the basic reproduction rate, *R*_0_(T), expressed as a function of temperature. Because absolute values of *R*_0_ depend on additional factors like host availability, breeding habitat quality, and vector control, we estimate a relative *R*_0_(T) (ie, scaled from 0 to 1) that isolates the impact of temperature on transmission, as in prior studies (25,71–73). Under this relative *R*_0_(T), the threshold condition *R*_0_(T) > 0 indicates the temperatures at which transmission is possible. This temperature-dependent transmission modeling approach has been found to reliably capture the temperature limits and peak of transmission in real-world settings (74).

We projected *R*_0_(T) using CA BCM temperature projections for MIROC under 4.5, 6.0, and 8.5 for the current, mid-century, and end-of-century time periods (as for the vector SDM; see Supplementary Methods). As the temperature input for *R*_0_(T), we specifically used the average of the maximum and minimum monthly temperature projections.

### Travel-associated cases

We estimated the number of travel-associated dengue cases occurring each month at the census tract level drawing upon four surveillance data sets: 1) annual county-level case reports and the putative region of exposure for 2016-2023 from the California Department of Public Health (CDPH) (14); 2) annual county-level case reports for 2010-2023 from the Centers for Disease Control and Prevention (75); 2) monthly, state-wide case reports for 2011-2023 from the CDPH (data not publicly available); and 3) annual, census tract-level travel-associated reported cases for Santa Clara and Los Angeles Counties from 2014-2024, obtained through data use agreements. Use of these data in this study was deemed non-human subjects research by the University of California, Berkeley ethics committee.

First, for each California census tract, we obtained population counts by racial and ethnic groups from the U.S. Census Bureau Detailed Demographic and Housing Characteristics File A (DHC-A) (76) and grouped these counts by world regions to align with CDPH data on regions of DENV exposure. Here, world regions align with the United Nations geographic region designations (UN Statistics Division, 2024): South Asia; Southeast Asia; East Asia; West Asia; Central America (including Mexico); South America; the Caribbean; Melanesia; Polynesia; as well as North, South, Middle, West, and East Africa (see Supplementary Table S1 for complete mapping) (10). We then multiplied these regionally-grouped counts by the proportion of travel-associated cases resulting from exposure in each region (Supplementary Table S1; CDPH, personal communication, March 4, 2024) to obtain an overall “risk of travel-associated cases” metric for each region. This approach therefore assumes that the population size of ethnic backgrounds from dengue endemic regions is correlated with the volume of travel to those regions.

We then modeled the number of travel-associated cases at the census-tract level, leveraging travel-associated case data from Santa Clara and Los Angeles Counties, as a function of these demographic risk metrics. We also included educational attainment, poverty, unemployment, and linguistic isolation as additional predictors (expressed as proportions of the census tract population and using data from the 2020 American Community Survey—ACS) (77), as these factors may influence international travel activity and impact the rates at which individuals can seek and access health care (78,79). These predictors were included in a Poisson generalized linear model estimating the number of travel-associated dengue cases using the total population size for a given census tract as a model offset (see Supplementary Methods for full model specification and further details about these predictors). We trained the model on total travel-associated dengue cases in census tracts in Santa Clara and Los Angeles Counties between 2014-2024, then predicted travel-associated cases over all census tracts in California. Lastly, to obtain monthly estimates, we multiplied predicted case counts by the proportion of cases reported per month across the state (Supplementary Methods). We summarized census tract-level predictions by county and found strong agreement with the actual number of reported cases at the county level (R^"^ = 0.94) (Supplementary Figure S2). However, to assess the sensitivity of our results to the sociodemographic assumptions described above, we also estimated the number of travel-associated cases at the county level using only the county-level case reports, multiplied by the average number of cases per month (ie, independent of county or tract-level sociodemographic data) (Supplementary Figure S3).

### Risk metric: Definition, calibration, and validation

We defined two related indices to quantify local dengue transmission risk: ‘ecological risk’, the product of vector presence and temperature-based suitability for pathogen transmission, and ‘eco-epidemiological risk’, the product of ecological risk and the number of predicted travel-associated cases (Figure 1). Both indices were calculated by raster multiplication of the input components.

To improve the interpretability of our risk metric in the context of real disease risk, we estimated risk for the instances of local transmission reported in California between 2023 and 2024 (eight unique locations and months). This included cases reported in Pasadena and Long Beach in October 2023; Baldwin Park and Panorama City in September 2024; Escondido, El Monte, and Hollywood Hills in October 2024; and the City of San Bernardino in November 2024 (Supplementary Table S2). We specifically estimated risk in the 1-2 months prior to when the case was reported given lags in transmission and case reporting and based on estimated symptom onset dates obtained through personal communication with city and/or state public health officials (CDPH, personal communication, March 12, 2025) (Supplementary Table S2). For each reported location/month of local transmission, we summarized census tract-level risk as the mean of ecological or eco-epidemiological risk, and used a spatial bootstrap (B = 1,000) that resampled pixels within each tract with probability proportional to population density. For reported locations spanning multiple census tracts, we included all intersecting tracts in our risk estimation. To define an empirical threshold for dengue risk, we then averaged the mean risk values from each instance of local transmission and obtained pooled 95% confidence intervals by repeatedly sampling one bootstrap draw from each instance and averaging across instances (pooled bootstrap, B = 5,000). To investigate the extent to which suitable conditions preceded observed local transmission, we also estimated monthly risk starting in 2016—by which point *Ae. aegypti* had been detected in at least 12 counties (57)—for each of the eight reported locations.

To assess transmission risk to residents of California, we estimated the ‘population at risk’, defined as the percent of the population living within 1 km (for ecological risk) or within a census tract (for eco-epidemiological risk) where risk exceeds that of observed local transmission. We estimated this percent using high-resolution population estimates (100 m grids) from ‘CA-POP’ (Supplementary Figure S4) (80).

Lastly, to validate our risk metric, we compared our estimates of ecological and eco-epidemiological risk during observed local transmission to those estimated at months and locations chosen at random (‘matched controls’) (n = 1,000). To ensure the randomly selected months and locations represented realistic transmission scenarios (eg, to avoid selecting unpopulated locations), census tracts were drawn with probabilities based on human population density, and months were drawn with probabilities based on the seasonality of travel to endemic regions (CDPH, personal communication, March 4, 2024) (Supplementary Figures S4-S5). We then assessed whether estimated risk was significantly higher in the locations and months of observed local transmission compared to the matched controls.

## Results

### Validation of risk metric

We developed two related risk indices—the first combining estimates of vector presence and temperature-dependent transmission suitability (‘ecological risk’), and the second incorporating an estimate of the number of travel-associated cases (‘eco-epidemiological risk’). Using these indices, we identified significantly higher risk for the locations and months in which local transmission was observed compared to matched controls for both ecological risk (t(7) = 4.25, p = 0.004) and eco-epidemiological risk (t(7) = 3.30, p = 0.013), suggesting that our metric can provide reliable estimates of heightened risk. In particular, our estimates of ecological and eco-epidemiological risk for the instances of observed local transmission were roughly three times higher than those for locations and times with similar population densities and seasonality of travel to endemic regions that did not report local transmission (means: 0.414 vs 0.140 for ecological risk and 0.0017 vs 0.0005 for eco-epidemiological risk; Supplementary Figure S5). However, only three individual instances of observed local transmission had a risk estimate significantly greater than expectations (eco-epidemiological risk for Pasadena in August 2023; and Baldwin Park and Panorama City in August 2024) (Supplementary Figure S5), suggesting that our metric is better suited to identify spatial and temporal patterns of risk rather than to serve as an operational warning tool.

When investigating risk at the locations across Los Angeles, San Bernardino, and San Diego Counties in which local dengue transmission was reported in 2023-2024 (Supplementary Table S2), we found that the risk was not uniquely high at the time of local transmission (Figure 2, Supplementary Figure S6). That is, for each of these locations, our ecological or eco-epidemiological risk estimate for the month of observed local transmission typically denoted a maximum for that year but was exceeded in several prior years (2-8 prior years starting in 2016, depending on the location).

**Figure 2.**
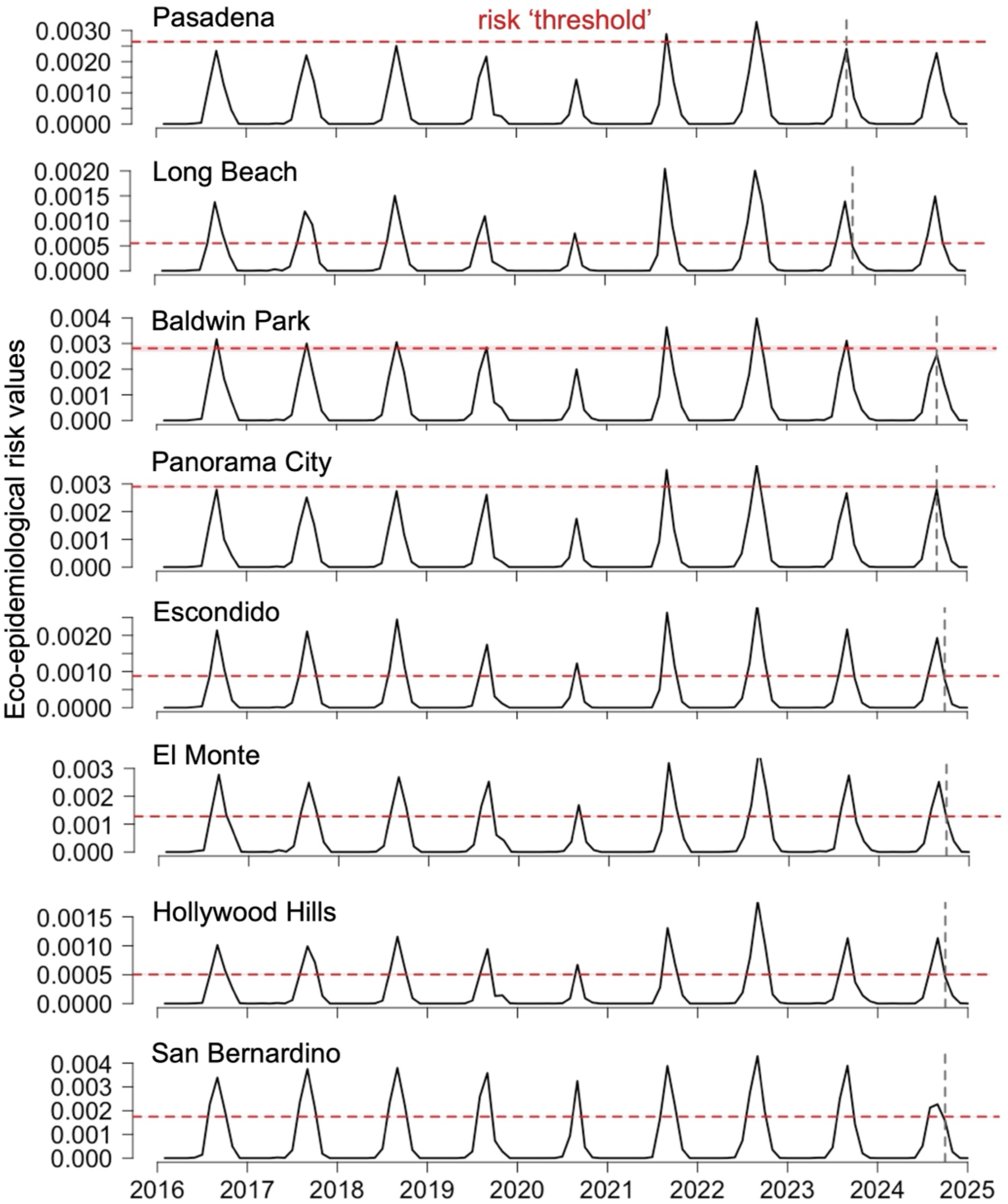
Monthly eco-epidemiological risk at locations of reported local transmission. Eco-epidemiological risk estimates (y-axis) are shown for each of the eight unique locations in which local dengue transmission was reported in California in 2023-2024. The gray, vertical dashed line in each plot denotes the timing of local transmission (estimated as 1-2 months prior to the case report). The red, horizontal dashed line and shading in each plot denote the mean and 95% confidence interval, respectively, of eco-epidemiological risk at the time of local transmission. See Supplementary Figure S6 for the corresponding time series of ecological risk estimates.

### Current patterns of risk and bottlenecks to local transmission

Under current environmental and sociodemographic conditions, we estimated relatively high suitability for dengue transmission in portions of the Central Valley and Los Angeles and San Diego metropolitan areas in July and August. That is, large portions of these regions experience eco-epidemiological risk at or above the ‘threshold’—the mean risk value estimated across instances of observed local transmission (Figure 3, Supplementary Figures S8-9). In particular, in the peak month of transmission risk (August), we estimated that approximately 18.2 million (95% CI: 17.9-18.3) California residents—46% of the state population—live in a region with eco-epidemiological risk at or above the threshold for observing local transmission (Figure 4, Supplementary Figure S9). Similarly, an estimated 17.6 million (95% CI: 17.2-18.1) (45%) residents currently live in a region above the threshold for ecological risk (Supplementary Figure S10).

**Figure 3.**
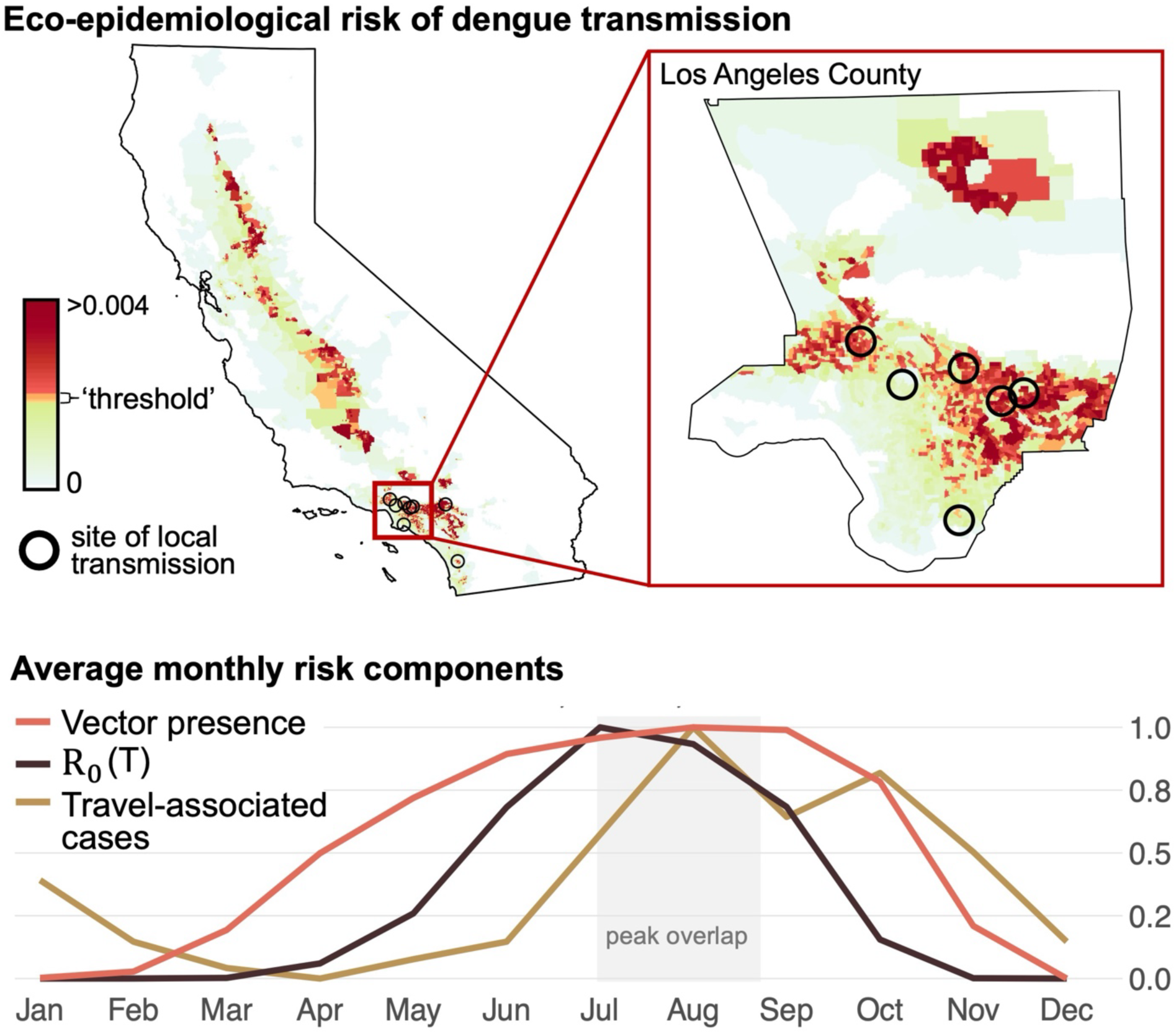
Current risk of dengue transmission across California. (Top) Estimates of eco-epidemiological risk of transmission in August (the month of peak risk) shown at the census tract level. White to green denotes 0 to low risk, yellow to orange denotes levels of risk estimated during observed local transmission (‘threshold’), while dark red denotes risk above this threshold. The inset plot denotes eco-epidemiological risk for Los Angeles County, wherein the majority of the observed locally-acquired cases were reported. (Bottom) Monthly state-wide averages for three major components of dengue transmission risk estimated in this study: vector presence, temperature-dependent transmission suitability (*R*_0_(T)), and travel-associated cases. Estimates are based on current conditions and values are scaled from 0 to 1 based on each components’ respective minimum and maximum. The light gray box denotes the peak overlap in risk components (July and August).

**Figure 4.**
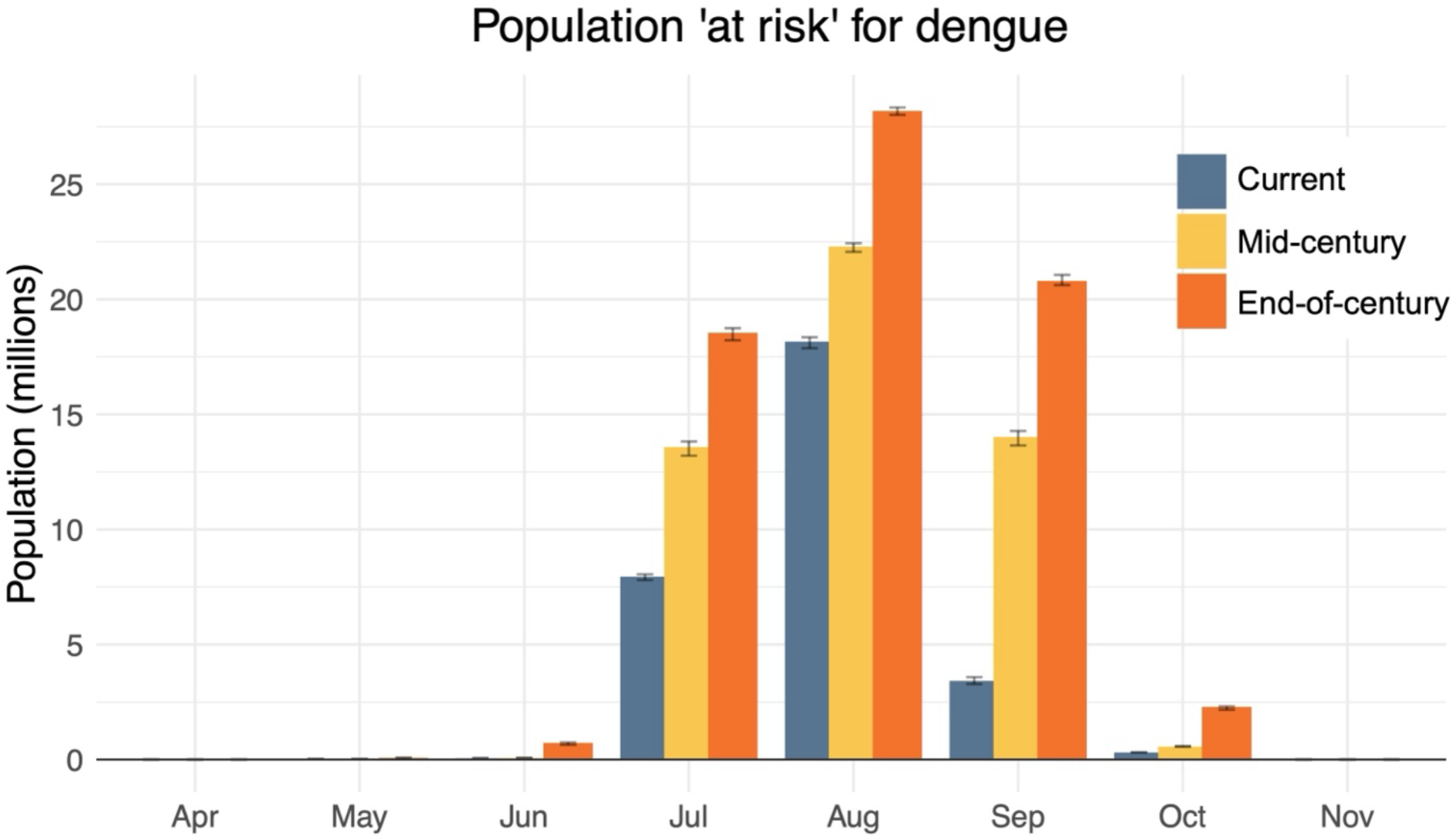
Population ‘at risk’ for dengue transmission. Bars denote the number of California residents (in millions) living within a census tract where the mean eco-epidemiological risk is at or above the ‘threshold.’ The threshold is based on the mean risk value estimated across census tracts and months with reported local transmission, while error bars denote estimates based on the lower and upper bounds of the 95% confidence interval around that mean. Estimates are shown for the current period (2010-2020), mid-century (2040-2050), and end-of-century (2090-2100) under a moderate climate warming scenario (RCP 4.5) (see Supplementary Figure S9 for estimates under RCP 6.0 and 8.5).

Outside populous portions of the Central Valley and Los Angeles and San Diego metropolitan areas, we estimated peak dengue risk to be at or near zero, as one or more of the required components for transmission is limiting (see Supplementary Figure S12 for regional bottlenecks to transmission). For example, for the western Central Valley, vector and temperature suitability may be sufficient for transmission in summer months, but infrequent travel-associated cases may limit risk. Low viral introductions, as well as low vector presence, may also limit transmission risk in the Inland desert region. In all regions of California, risk remains negligible outside June through October as temperatures are too low for transmission (Figure 3, Supplementary Figures S7-11, S14), suggesting that climate suitability is the major bottleneck to transmission outside of the summer months.

### Shifting patterns of risk under climate and land use change

We estimate that the ecological suitability for dengue transmission is generally expected to increase across California under projected climate warming and expansion of developed and urban land cover. In particular, we estimate an increase in the ecological risk during peak months (July-August) in coming decades, with the greatest increases predicted for the Bay Area, Central Valley, and coastal southern California regions (Figure 5, Supplementary Figure S15). We estimate that an additional 4.1 million (95% CI: 3.7-4.6) California residents will live in a region above the eco-epidemiological risk threshold by mid-century, and an additional 10.0 million (95% CI: 9.7-10.5) may be at risk by end-of-century (estimates are for August under a moderate climate warming scenario, RCP 4.5) (Figure 4) (see Supplementary Figure S9 for estimates under RCP 6.0 and 8.5). We note that these estimates are driven by projected shifts in vector presence and temperature-based suitability for transmission (Supplementary Figures S16-19), as we did not attempt to project changes in travel-associated cases or population growth. Based on ecological risk, we estimate that an additional 3.0 million (95% CI: 2.5-3.6) and 9.1 million (95% CI: 8.6-9.6) California residents will live in a region above the risk threshold by mid-century and end-of-century, respectively (Supplementary Figure S10). We estimate the largest increases in population at risk for September by mid-century and October by end-of-century, suggesting an expansion of the potential transmission season in coming decades (Figures 3-5, Supplementary Figures S9-11).

**Figure 5.**
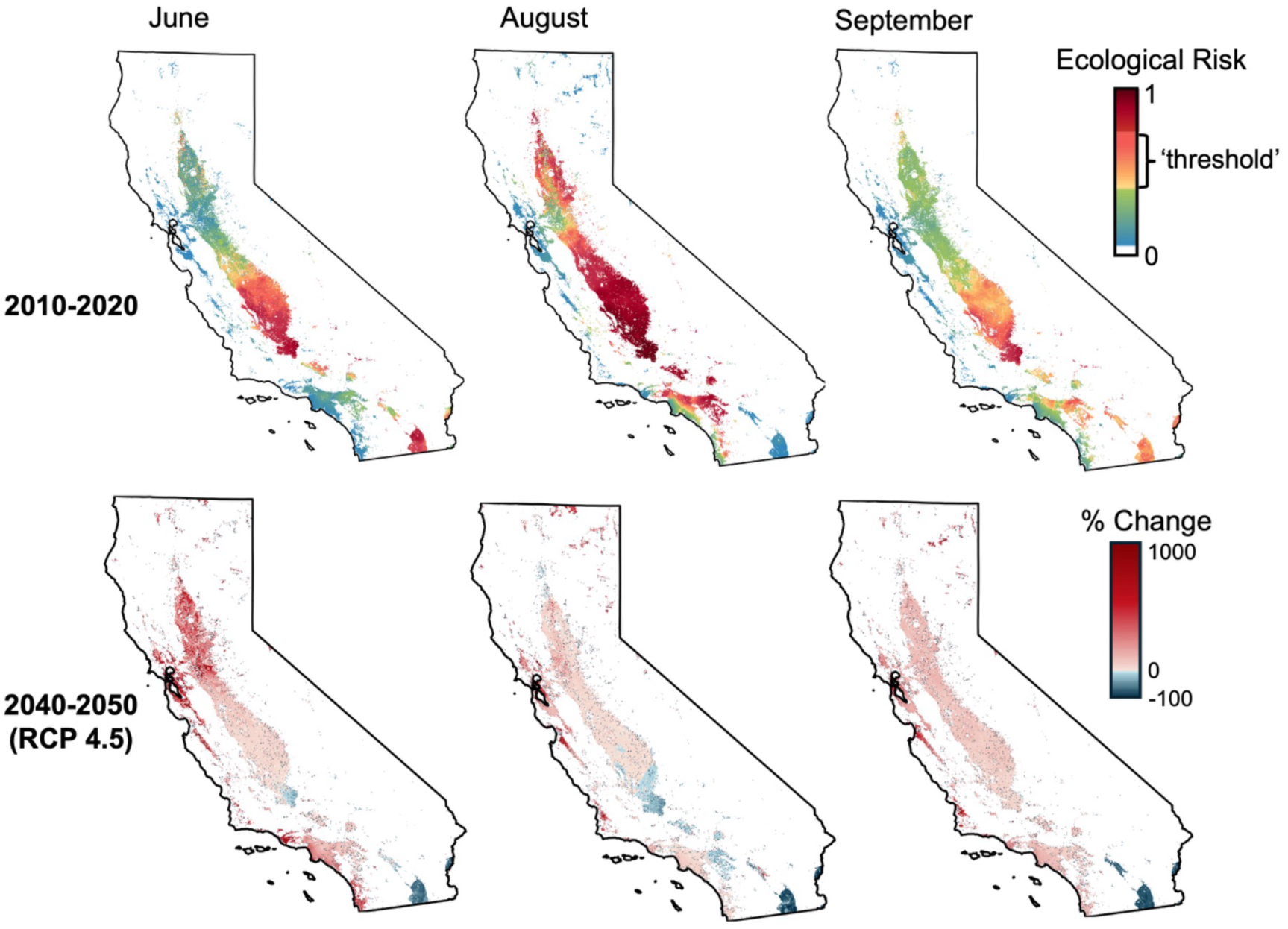
Shifting ecological risk of dengue transmission. Current (top panel) and mid-century (bottom panel) ecological risk of transmission. Estimates are shown for two shoulder months—June and September (left and right panels, respectively)—and one peak month—August (center panel). Colors in the top panel denote values of our estimated risk metric wherein white denotes zero risk, blue-green denotes low risk (ie, non-zero but below the risk ‘threshold’), and yellow-red denotes risk at or above the ‘threshold’ from observed local transmission. Colors in the bottom panel denote the percent change from current risk, calculated on a per-pixel (270 m) basis. See Supplementary Figure S7 for all months.

## Discussion

The rapid emergence of dengue in non-endemic regions poses a large and growing threat to public health, necessitating approaches to identify the regions and times at elevated risk. Here, we develop an approach to estimate dengue risk in emerging regions that integrates the key ecological and sociodemographic conditions needed for local transmission: vector presence, temperature suitability for pathogen transmission, and viral introductions via travel-associated cases. Applied to California—which reported its first locally-acquired dengue cases in 2023—our approach accurately identified the locations and months of observed local transmission as having elevated risk, suggesting it reliably captures spatial and temporal patterns of heightened risk.

Our results indicate that a substantial portion of the California population, including those in populous portions of the Central Valley and the Los Angeles and San Diego metropolitan areas, experience risk levels above those observed during local transmission in 2023-2024, which we used as a ‘threshold’ to aid interpretation in the context of real disease risk. Notably, we estimate that approximately 18.2 million (95% CI: 17.9-18.3) California residents (46%) currently live in areas where peak monthly risk exceeds this threshold. An additional 4.1 million (95% CI: 3.7-4.6) California residents may do so by mid-century under moderate projections of climate warming and urban expansion (ie, conversion to urban/developed land cover; population growth projections were not incorporated). Thus, while few locally-acquired cases have been reported to date (20 at the start of 2025), conditions in the state are likely suitable for additional outbreaks in coming years.

We found that local dengue transmission was likely possible in regions of southern California for several years prior to the observed instances in 2023-2024. That is, for the locations within Los Angeles, San Bernardino, and San Diego counties where local transmission was reported, we estimated higher risk levels in two or more prior years. In particular, suitable climate and land cover conditions were likely present in these locations prior to 2013 when breeding populations of *Aedes aegypti* were first detected in southern California (81). Despite aggressive vector control efforts, *Ae. aegypti* rapidly expanded across the state and was detected in at least 12 counties by 2016 (55,57). Thus, the first cases of locally-acquired dengue likely did not reflect a uniquely high overlap in ecological and sociodemographic conditions, but rather a combination of sufficient suitability and highly stochastic transmission dynamics typical of non-endemic regions with low viral circulation (82,83). This period between transmission suitability and disease emergence (eg, the ‘waiting time’) has been well described in the theoretical literature on critical transitions in disease states (84–86), wherein higher rates of travel-associated infections facilitates more sustained outbreaks of local transmission (32,54,87). As the global burden of dengue continues to rise, with over 14.1 million cases reported in 2024—the highest level recorded to date (8,10)—viral introductions to California may increase, potentially facilitating larger outbreaks.

In addition to estimating overall patterns of dengue risk, by separately estimating three of the key conditions necessary for transmission, our approach helps to identify major bottlenecks to transmission in different regions and times. For example, for populous regions such as the San Francisco Bay Area, dengue transmission is currently inhibited by low climate suitability as temperatures there are typically well below 29°C (the optimum for dengue transmission by *Ae. aegypti*) (25) throughout the year. Thus, climate warming could have a relatively large impact on risk in this region, as even moderate increases in temperature may relax thermal constraints on mosquito and pathogen life history traits, leading to large increases in transmission suitability—a pattern that has been found for West Nile virus transmission in California (27), and for dengue transmission more broadly (28). However, we note that, while we predict reasonable suitability for *Ae. aegypti* in this region, its actual presence may be substantially lower given extensive vector control efforts therein (57), suggesting that reduced intensity or effectiveness of vector control (eg, through the development of insecticide resistance) (88) may relax this constraint on transmission. Conversely, for less populous portions of the Central Valley, temperatures are suitable for transmission from roughly June to September, yet risk remains low as travel-associated cases are infrequent, suggesting that climate suitability is not the primary bottleneck to transmission in this region. However, we note that the true frequency of travel-associated cases may be substantially higher than what is reported, and under-ascertainment may vary spatially (89). Further, as the population size of California’s Central Valley is projected to increase in coming decades (90), as is global dengue incidence (8,10), viral introductions to this region may become more frequent, suggesting that enhanced public and physician awareness of dengue, early case detection, and active DENV surveillance in local *Aedes* populations may be warranted.

Our approach has several limitations that warrant consideration. First, our modeled estimates of travel-associated cases assume that the volume of travel to dengue endemic regions scales with the size of populations associated with those regions based on self-reported ethnicity, and that state-wide trends in the region of exposure apply to all census tracts—assumptions that likely oversimplify more complex travel dynamics. Further, although the seasonality of dengue transmission varies in different endemic regions from which travelers are returning (91–94) and may shift with climate change (95), we did not account for this in our estimate given limited data on travel by region and month. Similarly, we did not attempt to project changes in travel-associated cases, potentially leading us to underestimate future local transmission (8,21). We also did not incorporate potential variation in exposure to mosquitoes, which may be governed by factors such as housing quality and air conditioning usage (96), as comprehensive data on these fine-scale drivers of exposure are unavailable. However, we did incorporate poverty as a predictor when estimating travel-associated cases, which may partially account for socioeconomic heterogeneities in exposure risk (96). Additionally, as *Ae. aegypti* is still actively invading California, its distribution may be impacted by dispersal processes rather than driven solely by climate and land cover conditions, although we did attempt to control for this effect by excluding the first three years of surveillance data (ie, 2013-2015) when training our model.

Finally, while we attempt to provide an interpretable risk metric by defining a threshold risk value based on observed local transmission, this value does not represent a strict threshold (eg, no underlying physiological processes delineate values above and below this threshold) and is based on relatively few unique instances of local transmission (eight unique location-months) from case reporting, which may be subject to non-random underreporting. Given these limitations, we emphasize that our approach is best suited to estimate spatial and temporal patterns of risk rather than to serve as an operational early warning system for dengue in California.

While our analysis focuses on California as an emerging center of dengue risk in the continental USA, our finding that transmission suitability is relatively high in portions of the state likely indicates suitability for other regions in the southern USA and southern Europe. In particular, metropolitan areas in Arizona, Texas, and Florida and major European cities including Madrid and Paris have experienced locally-acquired dengue cases in recent years and may represent areas of increasing transmission risk given their overlap in *Aedes* vector presence, high population density, and suitable temperatures for pathogen transmission (97–99). Our approach of combining diverse data sources to estimate the underlying ecological, sociodemographic, and epidemiological conditions necessary for local transmission can be reasonably applied to these and other emerging regions. Estimating these underlying components separately enables identification of regionally-specific drivers and bottlenecks to transmission, which can inform targeted disease prevention strategies such as enhanced vector surveillance, surveillance of returning travelers, and public awareness campaigns. Our findings indicate that these preventative measures may become increasingly necessary in non-endemic regions under ongoing climate warming, urbanization, and global travel.

## Contributors

LIC and AJM originally conceived of the research project. All authors provided conceptual guidance and supported the project administration. LIC, TJS, SS, KPL, ZR, KMS, and AJM conducted formal analysis of the environmental, vector, and/or epidemiological data. LIC and AJM communicated with applied public health partners to obtain information on travel-associated cases and discuss preliminary results. LIC wrote the original draft. All authors received and edited the final draft and approved of submission.

## Data sharing

All R scripts used in the analysis and compiled data products are available in a publicly available GitHub repository: https://github.com/lcouper/CA-Dengue/. Raw vector surveillance and travel-associated case data were obtained through Data Use Agreements and cannot be directly shared.

## Declaration of interest

The authors have no competing interests to declare.

## Acknowledgements

LIC was funded by a National Science Foundation Postdoctoral Research Fellowship in Biology. AJM was funded by the USDA National Institute of Food and Agriculture (2023-68016-40683) and the National Science Foundation and Fogarty International Center (DEB-2011147; DEB-2339209). We gratefully acknowledge expert input from the California Department of Public Health, the City of Pasadena Public Health Department, the City of Long Beach Public Health Department, the Santa Clara County Public Health Department, and the Los Angeles County Public Health Department. We also acknowledge the California Vectorborne Diseases Surveillance System for providing the *Aedes aegypti* surveillance data used in this study.

## Notes

https://github.com/lcouper/CA-Dengue/

### Competing Interest Statement

The authors have declared no competing interest.

## References

1. Myers SS, Patz JA. Emerging Threats to Human Health from Global Environmental Change. Annual Review of Environment and Resources. 2009 Nov 21;34(Volume 34, 2009):223–52.

2. Malhi Y, Franklin J, Seddon N, Solan M, Turner MG, Field CB, et al. Climate change and ecosystems: threats, opportunities and solutions. Philosophical Transactions of the Royal Society B: Biological Sciences. 2020 Jan 27;375(1794):20190104.

3. Abbass K, Qasim MZ, Song H, Murshed M, Mahmood H, Younis I. A review of the global climate change impacts, adaptation, and sustainable mitigation measures. Environ Sci Pollut Res. 2022 June 1;29(28):42539–59.

4. Mahon MB, Sack A, Aleuy OA, Barbera C, Brown E, Buelow H, et al. A meta-analysis on global change drivers and the risk of infectious disease. Nature. 2024 May 23;629(8013):830–6.

5. Van de Vuurst P, Escobar LE. Climate change and infectious disease: a review of evidence and research trends. Infect Dis Poverty. 2023 May 16;12(1):51.

6. Caminade C, McIntyre KM, Jones AE. Impact of recent and future climate change on vector-borne diseases. Annals of the New York Academy of Sciences. 2019;1436(1):157–73.

7. Sah R, Siddiq A, Padhi BK, Mohanty A, Rabaan AA, Chandran D, et al. Dengue virus and its recent outbreaks: current scenario and counteracting strategies. Int J Surg. 2023 Mar 1;109(9):2841–5.

8. Messina JP, Brady OJ, Golding N, Kraemer MUG, Wint GRW, Ray SE, et al. The current and future global distribution and population at risk of dengue. Nature Microbiology. 2019 Sept;4(9):1508–15.

9. Harish V, Colón-González FJ, Moreira FRR, Gibb R, Kraemer MUG, Davis M, et al. Human movement and environmental barriers shape the emergence of dengue. Nat Commun. 2024 May 28;15(1):4205.

10. World Health Organization. Dengue - global situtation [Internet]. 2023. (Disease Outbreak News). Available from: https://www.who.int/emergencies/disease-outbreak-news/item/2023-DON498

11. Ryan SJ, Carlson CJ, Mordecai EA, Johnson LR. Global expansion and redistribution of Aedes-borne virus transmission risk with climate change. PLOS Neglected Tropical Diseases. 2019 Mar 28;13(3):e0007213.

12. Murray NEA, Quam M, Wilder-Smith A. Epidemiology of dengue: past, present and future prospects. CLEP. 2013 Aug;299.

13. CDC. Centers for Disease Control and Prevention. 2024 [cited 2024 Feb 12]. Dengue areas of risk in the US | CDC. Available from: https://www.cdc.gov/dengue/areaswithrisk/in-the-us.html

14. CDPH. CDPH Monthly Update on Number of Dengue Infections in California February 1, 2024. California Department of Public Health; 2024.

15. Wong JM, Rivera A, Volkman HR, Torres-Velasquez B, Rodriguez DM, Paz-Bailey G, et al. Travel-Associated Dengue Cases — United States, 2010–2021. 2023;72(30).

16. Pan American Health Organization / World Heatlh Organization. Epidemiological Update: Increase in dengue cases in the Region of the Americas. Washington D.C.; 2024.

17. Robert MA, Christofferson RC, Weber PD, Wearing HJ. Temperature impacts on dengue emergence in the United States: Investigating the role of seasonality and climate change. Epidemics. 2019 Sept 1;28:100344.

18. Brady OJ, Hay SI. The Global Expansion of Dengue: How Aedes aegypti Mosquitoes Enabled the First Pandemic Arbovirus. Annual Review of Entomology. 2020 Jan 7;65(Volume 65, 2020):191–208.

19. Kamal M, Kenawy MA, Rady MH, Khaled AS, Samy AM. Mapping the global potential distributions of two arboviral vectors Aedes aegypti and Ae. albopictus under changing climate. PLOS ONE. 2018 Dec 31;13(12):e0210122.

20. Dupuis B, Brézillon-Dubus L, Failloux AB. [The effects of climate change on the emergence of dengue]. Med Sci (Paris). 2025 Feb;41(2):137–44.

21. Childs ML, Lyberger K, Harris M, Burke M, Mordecai EA. Climate warming is expanding dengue burden in the Americas and Asia [Internet]. 2024 [cited 2025 May 3]. Available from: http://medrxiv.org/lookup/doi/10.1101/2024.01.08.24301015

22. Kamiya T, Greischar MA, Wadhawan K, Gilbert B, Paaijmans K, Mideo N. Temperature-dependent variation in the extrinsic incubation period elevates the risk of vector-borne disease emergence. Epidemics. 2020 Mar 1;30:100382.

23. Mordecai EA, Caldwell JM, Grossman MK, Lippi CA, Johnson LR, Neira M, et al. Thermal biology of mosquito-borne disease. Ecology Letters. 2019;22(10):1690–708.

24. Parham P, Michael E. Modeling the Effects of Weather and Climate Change on Malaria Transmission. Environmental Health Perspectives. 2010 May 1;118(5):620–6.

25. Mordecai EA, Cohen JM, Evans MV, Gudapati P, Johnson LR, Lippi CA, et al. Detecting the impact of temperature on transmission of Zika, dengue, and chikungunya using mechanistic models. PLOS Neglected Tropical Diseases. 2017 Apr 27;11(4):e0005568.

26. Liu-Helmersson J, Stenlund H, Wilder-Smith A, Rocklöv J. Vectorial Capacity of Aedes aegypti: Effects of Temperature and Implications for Global Dengue Epidemic Potential. PLOS ONE. 2014 Mar 6;9(3):e89783.

27. Skaff NK, Cheng Q, Clemesha RES, Collender PA, Gershunov A, Head JR, et al. Thermal thresholds heighten sensitivity of West Nile virus transmission to changing temperatures in coastal California. Proc R Soc B. 2020 Aug 12;287(1932):20201065.

28. Caldwell JM, LaBeaud AD, Lambin EF, Stewart-Ibarra AM, Ndenga BA, Mutuku FM, et al. Climate predicts geographic and temporal variation in mosquito-borne disease dynamics on two continents. Nat Commun. 2021 Feb 23;12(1):1233.

29. Peña-García VH, Triana-Chávez O, Arboleda-Sánchez S. Estimating Effects of Temperature on Dengue Transmission in Colombian Cities. Annals of Global Health. 2017 Nov 21;83(3–4):509.

30. Wilder-Smith A, Gubler DJ. Geographic Expansion of Dengue: The Impact of International Travel. Medical Clinics of North America. 2008 Nov 1;92(6):1377–90.

31. World Health Organization. Dengue: Guidelines for Diagnosis, Treatment, Prevention and Control. World Health Organization; 2009. 159 p.

32. Gwee XWS. Global dengue importation: a systematic review. 2021;

33. Tatem AJ, Hay SI, Rogers DJ. Global traffic and disease vector dispersal. Proceedings of the National Academy of Sciences. 2006 Apr 18;103(16):6242–7.

34. Powell JR, Gloria-Soria A, Kotsakiozi P. Recent History of Aedes aegypti: Vector Genomics and Epidemiology Records. BioScience. 2018 Nov 1;68(11):854–60.

35. Powell JR, Tabachnick WJ. History of domestication and spread of Aedes aegypti - A Review. Mem Inst Oswaldo Cruz. 2013 Dec;108(Suppl 1):11–7.

36. Reiter P, Amador MA, Anderson RA, Clark GG. Short report: dispersal of Aedes aegypti in an urban area after blood feeding as demonstrated by rubidium-marked eggs. Am J Trop Med Hyg. 1995 Feb;52(2):177–9.

37. Cornel AJ, Hunt RH. Aedes albopictus in Africa? First records of live specimens in imported tires in Cape Town. J Am Mosq Control Assoc. 1991 Mar 1;7(1):107–8.

38. Adhami J, Reiter P. Introduction and establishment of Aedes (Stegomyia) albopictus skuse (Diptera: Culicidae) in Albania. J Am Mosq Control Assoc. 1998 Sept 1;14(3):340–3.

39. Hawley WA, Reiter P, Copeland RS, Pumpuni CB, Craig GB. Aedes albopictus in North America: Probable Introduction in Used Tires from Northern Asia. Science. 1987 May 29;236(4805):1114–6.

40. Romi R. History and updating on the spread of Aedes albopictus in Italy. Parassitologia. 1995 Dec 1;37(2–3):99–103.

41. Swan T, Russell TL, Staunton KM, Field MA, Ritchie SA, Burkot TR. A literature review of dispersal pathways of Aedes albopictus across different spatial scales: implications for vector surveillance. Parasites & Vectors. 2022 Aug 27;15(1):303.

42. Simmons CP, Farrar JJ, Vinh Chau, Nguyen van, Bridget W. Dengue. New England Journal of Medicine. 2012;366(15):1423–32.

43. Vulu F, Futami K, Sunahara T, Mampuya P, Bobanga TL, Mumba Ngoyi D, et al. Geographic expansion of the introduced Aedes albopictus and other native Aedes species in the Democratic Republic of the Congo. Parasites & Vectors. 2024 Jan 26;17(1):35.

44. Armbruster PA. Photoperiodic Diapause and the Establishment of Aedes albopictus (Diptera: Culicidae) in North America. Journal of Medical Entomology. 2016 Sept 1;53(5):1013–23.

45. Kraemer MU, Sinka ME, Duda KA, Mylne AQ, Shearer FM, Barker CM, et al. The global distribution of the arbovirus vectors Aedes aegypti and Ae. albopictus. eLife [Internet]. 2015 June 30 [cited 2020 May 8];4. Available from: https://elifesciences.org/articles/08347

46. Ngoagouni C, Kamgang B, Nakouné E, Paupy C, Kazanji M. Invasion of Aedes albopictus (Diptera: Culicidae) into central Africa: what consequences for emerging diseases? Parasites Vectors. 2015 Mar 31;8(1):191.

47. Ratnam I, Leder K, Black J, Torresi J. Dengue fever and international travel. J Travel Med. 2013 Dec;20(6):384–93.

48. Gubler DJ. Dengue and Dengue Hemorrhagic Fever. CLIN MICROBIOL REV. 1998;11:17.

49. Schwartz E, Weld LH, Wilder-Smith A, von Sonnenburg F, Keystone JS, Kain KC, et al. Seasonality, Annual Trends, and Characteristics of Dengue among Ill Returned Travelers, 1997–2006. Emerg Infect Dis. 2008 July;14(7):1081–8.

50. Tatem AJ, Huang Z, Das A, Qi Q, Roth J, Qiu Y. Air travel and vector-borne disease movement. Parasitology. 2012 Dec;139(14):1816–30.

51. Frank C, Schöneberg I, Krause G, Claus H, Ammon A, Stark K. Increase in Imported Dengue, Germany, 2001–2002. Emerg Infect Dis. 2004 May;10(5):903–6.

52. Wilder-Smith A. Dengue infections in travellers. Paediatr Int Child Health. 2012 May;32(s1):28–32.

53. Allwinn R. Significant increase in travel-associated dengue fever in Germany. Med Microbiol Immunol. 2011 Aug 1;200(3):155–9.

54. Poongavanan J, Lourenço J, Tsui JLH, Colizza V, Ramphal Y, Baxter C, et al. Dengue virus importation risks in Africa: a modelling study. The Lancet Planetary Health. 2024 Dec 1;8(12):e1043–54.

55. Pless E, Gloria-Soria A, Evans BR, Kramer V, Bolling BG, Tabachnick WJ, et al. Multiple introductions of the dengue vector, Aedes aegypti, into California. PLOS Neglected Tropical Diseases. 2017 Aug 10;11(8):e0005718.

56. Lee Y, Schmidt H, Collier TC, Conner WR, Hanemaaijer MJ, Slatkin M, et al. Genome-wide divergence among invasive populations of Aedes aegypti in California. BMC Genomics. 2019 Mar 12;20(1):204.

57. Metzger ME, Hardstone Yoshimizu M, Padgett KA, Hu R, Kramer VL. Detection and Establishment of Aedes aegypti and Aedes albopictus (Diptera: Culicidae) Mosquitoes in California, 2011–2015. Journal of Medical Entomology. 2017 May 1;54(3):533–43.

58. Shankar MB, Rodríguez-Acosta RL, Sharp TM, Tomashek KM, Margolis HS, Meltzer MI. Estimating dengue under-reporting in Puerto Rico using a multiplier model. PLOS Neglected Tropical Diseases. 2018 Aug 6;12(8):e0006650.

59. Chen T, Guestrin C. XGBoost: A Scalable Tree Boosting System [Internet]. 2016 [cited 2022 June 1]. Available from: https://dl.acm.org/doi/10.1145/2939672.2939785

60. Madon MB, Mulla MS, Shaw MW, Kluh S, Hazelrigg JE. Introduction of Aedes albopictus (Skuse) in southern California and potential for its establishment. J Vector Ecol. 2002 June 1;27(1):149–54.

61. Linthicum K, Vl K, Mb M, K F, undefined. Introduction and potential establishment of Aedes albopictus in California in 2001. J Am Mosq Control Assoc. 2003 Dec 1;19(4):301–8.

62. Gloria-Soria A, Payne AF, Bialosuknia SM, Stout J, Mathias N, Eastwood G, et al. Vector Competence of Aedes albopictus Populations from the Northeastern United States for Chikungunya, Dengue, and Zika Viruses. Am J Trop Med Hyg. 2021 Mar;104(3):1123–30.

63. Porse CC, Kramer V, Yoshimizu MH, Metzger M, Hu R, Padgett K, et al. Public Health Response to Aedes aegypti and Ae. albopictus Mosquitoes Invading California, USA. Emerg Infect Dis. 2015 Oct;21(10):1827–9.

64. Lambrechts L, Scott TW, Gubler DJ. Consequences of the Expanding Global Distribution of Aedes albopictus for Dengue Virus Transmission. PLOS Neglected Tropical Diseases. 2010 May 25;4(5):e646.

65. Flint LE, Flint AL, Thorne JH, Boynton R. Fine-scale hydrologic modeling for regional landscape applications: the California Basin Characterization Model development and performance. Ecol Process. 2013 Dec;2(1):25.

66. Rose NH, Badolo A, Sylla M, Akorli J, Otoo S, Gloria-Soria A, et al. Dating the origin and spread of specialization on human hosts in Aedes aegypti mosquitoes. eLife. 2023 Mar 10;12:e83524.

67. Oeser J, Zurell D, Mayer F, Çoraman E, Toshkova N, Deleva S, et al. The Best of Two Worlds: Using Stacked Generalisation for Integrating Expert Range Maps in Species Distribution Models. Global Ecology and Biogeography. 2024;n/a(n/a):e13911.

68. Valavi R, Guillera-Arroita G, Lahoz-Monfort JJ, Elith J. Predictive performance of presence-only species distribution models: a benchmark study with reproducible code. Ecological Monographs. 2022;92(1):e01486.

69. Araújo MB, New M. Ensemble forecasting of species distributions. Trends in Ecology & Evolution. 2007 Jan 1;22(1):42–7.

70. Sohl TL, Sayler KL, Bouchard MA, Reker RR, Friesz AM, Bennett SL, et al. Spatially explicit modeling of 1992–2100 land cover and forest stand age for the conterminous United States. Ecological Applications. 2014;24(5):1015–36.

71. Ryan SJ, McNally A, Johnson LR, Mordecai EA, Ben-Horin T, Paaijmans K, et al. Mapping Physiological Suitability Limits for Malaria in Africa Under Climate Change. Vector-Borne and Zoonotic Diseases. 2015 Nov 18;15(12):718–25.

72. Heidecke J, Wallin J, Fransson P, Singh P, Sjödin H, Stiles PC, et al. Uncovering temperature sensitivity of West Nile virus transmission: Novel computational approaches to mosquito-pathogen trait responses. Ten Bosch Q, editor. PLoS Comput Biol. 2025 Mar 31;21(3):e1012866.

73. Shocket MS, Verwillow AB, Numazu MG, Slamani H, Cohen JM, El Moustaid F, et al. Transmission of West Nile and five other temperate mosquito-borne viruses peaks at temperatures between 23°C and 26°C. eLife. 2020 Sept 15;9:e58511.

74. Mordecai EA, Ryan SJ, Caldwell JM, Shah MM, LaBeaud AD. Climate change could shift disease burden from malaria to arboviruses in Africa. The Lancet Planetary Health. 2020 Sept;4(9):e416–23.

75. Centers for Disease Control and Prevention. Dengue Historic Data (2010-2014) [Internet]. 2025. Available from: https://www.cdc.gov/dengue/data-research/facts-stats/historic-data.html

76. U.S. Census Bureau. 2020 Census Detailed Demographic and Housing Characteristics File A (Detailed DHC-A). 2020.

77. U.S. Census Bureau. 2020 American Community Survey. 2020.

78. Carabali M, Hernandez LM, Arauz MJ, Villar LA, Ridde V. Why are people with dengue dying? A scoping review of determinants for dengue mortality. BMC Infect Dis [Internet]. 2015 Dec [cited 2025 May 1];15(1). Available from: http://bmcinfectdis.biomedcentral.com/articles/10.1186/s12879-015-1058-x

79. Ng TC, Teo CH, Toh JY, Dunn AG, Ng CJ, Ang TF, et al. Factors influencing healthcare seeking in patients with dengue: Systematic review. Tropical Med Int Health. 2022 Jan;27(1):13–27.

80. Depsky NJ, Cushing L, Morello-Frosch R. High-resolution gridded estimates of population sociodemographics from the 2020 census in California. Vadrevu KP, editor. PLoS ONE. 2022 July 14;17(7):e0270746.

81. Gloria-Soria A, Brown JE, Kramer V, Hardstone Yoshimizu M, Powell JR. Origin of the Dengue Fever Mosquito, Aedes aegypti, in California. Sang RC, editor. PLoS Negl Trop Dis. 2014 July 31;8(7):e3029.

82. Morens DM, Folkers GK, Fauci AS. The challenge of emerging and re-emerging infectious diseases. Nature. 2004 July 8;430(6996):242–9.

83. Stollenwerk N, Mateus L, Steindorf V, Guerrero BV, Blasco-Aguado R, Cevidanes A, et al. Evaluating the risk of mosquito-borne diseases in non-endemic regions: A dynamic modeling approach [Internet]. medRxiv; 2024 [cited 2025 Apr 9]. p. 2024.10.10.24315163. Available from: https://www.medrxiv.org/content/10.1101/2024.10.10.24315163v1

84. Dibble CJ, O’Dea EB, Park AW, Drake JM. Waiting time to infectious disease emergence. Journal of The Royal Society Interface [Internet]. 2016 Oct 31 [cited 2025 Apr 9]; Available from: https://royalsocietypublishing.org/doi/10.1098/rsif.2016.0540

85. Czuppon P, Schertzer E, Blanquart F, Débarre F. The stochastic dynamics of early epidemics: probability of establishment, initial growth rate, and infection cluster size at first detection. Journal of The Royal Society Interface. 2021 Nov 17;18(184):20210575.

86. Drake JM, Brett TS, Chen S, Epureanu BI, Ferrari MJ, Marty É, et al. The statistics of epidemic transitions. Regoes RR, editor. PLoS Comput Biol. 2019 May 8;15(5):e1006917.

87. Yang X, Quam MBM, Zhang T, Sang S. Global burden for dengue and the evolving pattern in the past 30 years. Journal of Travel Medicine. 2021 Dec 29;28(8):taab146.

88. Yang F, Schildhauer S, Billeter SA, Hardstone Yoshimizu M, Payne R, Pakingan MJ, et al. Insecticide Resistance Status of Aedes aegypti (Diptera: Culicidae) in California by Biochemical Assays. Hribar L, editor. Journal of Medical Entomology. 2020 July 4;57(4):1176–83.

89. Gibbons CL, Mangen MJJ, Plass D, Havelaar AH, Brooke RJ, Kramarz P, et al. Measuring underreporting and under-ascertainment in infectious disease datasets: a comparison of methods. BMC Public Health. 2014 Dec;14(1):147.

90. California Department of Finance. Demographic Research Unit. State And County Population Projections 2020-2070. Sacramento, California: State of California Department of Finance; 2025.

91. Urmi TJ, Mosharrafa RA, Hossain MdJ, Rahman MS, Kadir MF, Islam MdR. Frequent outbreaks of dengue fever in South Asian countries—A correspondence analyzing causative factors and ways to avert. Health Sci Rep. 2023 Sept 28;6(10):e1598.

92. Stolerman LM, Maia PD, Kutz JN. Forecasting dengue fever in Brazil: An assessment of climate conditions. PLoS One. 2019 Aug 8;14(8):e0220106.

93. Carreto C, Gutiérrez-Romero R, Rodríguez T. Climate-driven mosquito-borne viral suitability index: measuring risk transmission of dengue, chikungunya and Zika in Mexico. International Journal of Health Geographics. 2022 Oct 27;21(1):15.

94. Ouattara CA, Traore S, Sangare I, Traore TI, Meda ZC, Savadogo LGB. Spatiotemporal analysis of dengue fever in Burkina Faso from 2016 to 2019. BMC Public Health. 2022 Mar 8;22(1):462.

95. Ebi KL, Nealon J. Dengue in a changing climate. Environmental Research. 2016 Nov 1;151:115–23.

96. Donnelly MAP, Kluh S, Snyder RE, Barker CM. Quantifying sociodemographic heterogeneities in the distribution of Aedes aegypti among California households. Viennet E, editor. PLoS Negl Trop Dis. 2020 July 21;14(7):e0008408.

97. Centers for Disease Control and Prevention. Data and Statistics on Dengue in the United States. 2024.

98. Redondo-Bravo L, Ruiz-Huerta C, Gomez-Barroso D, Sierra-Moros MJ, Benito A, Herrador Z. Imported dengue in Spain: a nationwide analysis with predictive time series analyses. Journal of Travel Medicine. 2019 Dec 23;26(8):taz072.

99. Fournet N, Voiry N, Rozenberg J, Bassi C, Cassonnet C, Karch A, et al. A cluster of autochthonous dengue transmission in the Paris region – detection, epidemiology and control measures, France, October 2023. Eurosurveillance [Internet]. 2023 Dec 7 [cited 2025 May 28];28(49). Available from: https://www.eurosurveillance.org/content/10.2807/1560-7917.ES.2023.28.49.2300641

