## Supplementary Material for "Climate warming and urbanization may expand dengue transmission risk in California"

**Supplementary Methods**


1. Vector species distribution model
2. Travel-associated cases
3. Risk metric


**Supplementary Figures S1-S19**


**Fig S1.** Land cover classes
**Fig S2.** Predicted vs observed county-level dengue cases
**Fig S3.** Monthly county-level travel-associated case estimates
**Fig S4.** California population density estimates
**Fig S5.** Risk metric validation
**Fig S6.** Time series of ecological risk at locations of local transmission
**Fig S7.** Monthly estimates of current ecological risk
**Fig S8.** Monthly estimates of current eco-epidemiological risk
**Fig S9**. Population at eco-epidemiological risk for dengue transmission
**Fig S10**. Population at ecological risk for dengue transmission
**Fig S11.** Population at eco-epidemiological risk using county-level case estimates
**Fig S12.** Bottlenecks to transmission
**Fig S13**. Monthly values of risk and its constitutive components
**Fig S14.** State-wide average monthly temperatures
**Fig S15**. Monthly percent change in ecological risk
**Fig S16**. Monthly estimates of $R_0(T)$ for mid-century
**Fig S17.** Monthly estimates of $R_0(T)$ for end-of-century
**Fig S18**. Monthly estimates of vector presence for mid-century
**Fig S19.** Monthly estimates of vector presence for end-of-century


**Supplementary Tables S1-S3**


**Table S1.** Percent cases by region of exposure and mapping to race/ethnicity categories
**Table S2.** Risk estimates during observed local transmission
**Table S3.** Coefficient estimates from model estimating tract-level cases


**Supplementary References**

**Supplementary Methods**

1. *Vector species distribution model*

To estimate *Aedes aegypti* presence across California, we used vector occurrence data (described in the main text) within two common species distribution modeling approaches—Maximum Entropy (MaxEnt) and Extreme Gradient Boosting (here, XGBoost)—that have different underlying objectives, assumptions, and strengths. MaxEnt uses a 'maximum entropy' approach to relate species occurrences to environmental covariates (1). We implemented this approach using the ENMeval package in R (2), using the "LQPH" configuration for feature classes to enable flexible relationships between response and predictors, and using a regularization parameter of 1 to minimize overfitting when extrapolating to future projections. Gradient-boosted decision trees is a supervised machine learning approach wherein decision trees, which partition the predictors based on similarities among the response variable, are iteratively built from the prediction errors of the prior tree (3). This approach allows for complex, non-linear relationships between predictors and response, and the XGBoost implementation used here has been found to minimize overfitting and optimize predictive accuracy (4). Here, we tuned hyperparameters including tree depth, learning rate, the number of randomly selected predictors at each split, minimal node size, and minimum loss reduction through spatiotemporal cross-validation.

To train and validate the models, we generated background points for *Ae. aegypti* (n = 10,000) using a two-stage approach. In the first stage, we used monthly occurrences of adult *Culex quinquefasciatus* from the CalSurv trap surveillance data as this indicates where mosquito surveillance is occurring but *Ae. aegypti* was not identified in a given month. More specifically, we identified all *Ae. aegypti* and *Cx. quinquefasciatus* occurrences in a given month and year. We then thinned the *Ae. aegypti* points, retaining a single occurrence within a given 275 m buffer (given the 270 m spatial resolution of the climate predictors used in the model). We repeated this thinning process for *Cx. quinquefasciatus* and further removed any points within an *Ae. aegypti* buffer to maintain priority of the *Ae. aegypti* observations. Next, we used a temperature suitability mask to randomly sample background points, weighted toward regions known to be unsuitable based on thermal constraints to *Ae. aegypt*i population persistence (following methods in Kraemer et al. 2015) (5). That is, we sampled 90% of our background points outside of the temperature suitability range, and 10% within to reduce bias in variable importance towards temperature. For each month, we targeted inclusion of 1,000 more background points than presence points after thinning (eg, 1,037 background points for 37 *Ae. aegypti* presence points). Background points were then randomly assigned to a month-year using weights proportional to the distribution of month-years from the presence points.

For both modeling approaches, our environmental covariates included mean temperature, total precipitation, and terrestrial water storage as predictors (see main text). We initially considered climatic water deficit, but it was highly collinear with mean temperature and was thus dropped. Further, we initially included land cover classes as a predictor in the SDMs. However, this resulted in high vector presence probability predicted in parts of California with temperature suitability for *Ae. aegypti,* but with land cover types unlikely to harbor this species based on prior studies such as grasslands or shrublands (5–7), suggesting that the model was not sufficiently sensitive to land cover. Thus, we instead masked outputs from both modeling approaches based on land cover types that are highly unlikely to harbor *Ae. aegypti*, rather than including land cover in models directly. This included masking all land cover types except those associated with high human activity, including developed, cropland, and hay/pastureland (ie, removing water, mining, barren, forest, grassland, shrubland, wetlands, ice/snow, and mechanically disturbed public/private lands).

Lastly, results from the two SDMs were used as input into a meta-learner to obtain final estimates of vector presence probability, ensuring our results were robust to modeling approaches and assumptions. Our meta-learner used a random forest algorithm tuned through spatiotemporal cross-validation and performed slightly better than either individual learner alone.

We note that our SDM does not account for vector control measures, which have been extensive since *Ae. aegypti* was detected in 2013 and may vary year-to-year, seasonally, and by region (8). Although control measures would alter the true monthly distribution of vectors from that expected due to climate and land cover features, comprehensive data on their timing and location are not yet available. However, our SDM uses presence-absence data, which may be less influenced by vector control than abundance. For example, the identification of a single *Ae. aegypti* in a county may elicit aggressive vector control leading to low abundance and reduced spread, but that individual would still appear as a presence in the surveillance record.

2. *Travel-associated cases*

To estimate census tract-level travel-associated cases, we first grouped population counts of reported racial and ethnic groups from the U.S. Census, 2020 Detailed Demographic and Housing Characteristics File A (9), based on world regions: South Asia, Southeast Asia, East Asia, West Asia, Central America (including Mexico), South America, the Caribbean, Melanesia, Polynesia, as well as North, South, Middle, West and East Africa (Supplementary Table S1). We then multiplied these regionally-grouped counts by the proportion of travel-associated cases from each region based on state-wide trends identified by the California Department of Public Health (Supplementary Table S1) to obtain a 'demographic risk of travel-associated cases' metric for each demographic group/region in each census tract, t (eg, 0.49*Population from Central America$_t$, 0.25*Population from South Asia$_t$, etc.).

Next, we modeled the number of travel-associated cases for a given tract ($C_t$) in Los Angeles and Santa Clara Counties—two of the top three counties in number of travel-associated dengue cases in California, and that fall within distinct regions of the state—using a Poisson generalized linear model. In the full model, we include the demographic risk metrics calculated above for the top three regions contributing travel-associated cases (South Asia: $SA_t$, South East Asia: $SEA_t$, and Central America: $CA_t$), as well as the total population within each census tract from racial and ethnic groups associated with any dengue endemic region, $DenPop_t$. Because travel-associated cases in different regions of California (e.g., Southern California, Bay Area) may predominantly come from exposure in different dengue endemic regions, we interact our demographic risk metrics with an indicator variable, $SoCal_t$, that takes the value 1 for census tracts in Southern California (eg, Los Angeles County), and the value 0 if the census tract is in Northern California (eg, Santa Clara County). We additionally include several sociodemographic predictors that may impact residents' ability to seek out and access health care, as well as the likelihood of international travel. These included educational attainment ($E_t$) (the percent of the population over 25 with less than a college education), poverty ($P_t$) (the percent of the population whose income in the past 12 months is below the federal poverty level), linguistic isolation ($L_t$) (the percent of households where no one over the age of 14 speaks English well), and unemployment ($U_t$) (the proportion of people over 16 who do not have a job but are available to work and have actively looked for work within the past month). Census tract-level data on these predictors were obtained from the American Community Survey (ACS). Each of these variables was standardized (z-score transformed) to assess relative effect sizes. Finally, we used the total population size for a given census tract ($TotPop_t$) as a model offset. The model specification is as follows:

$$C_t \sim Poisson(\lambda_t) \text{ with } log(\lambda_t) = \beta_0 + \beta_1\, Regions_t * SoCal_t + \beta_2\, DenPop_t + \beta_3\, Dems_t + log(TotPop_t)$$

where $Regions_t$ is a vector of the demographic risk metrics for the top three regions contributing travel-associated cases ($SA_t$, $SEA_t$, and $CA_t$), and $Dems_t$ is a vector of the four sociodemographic characteristics described above ($E_t$, $P_t$, $L_t$, and $U_t$). From this full model, we selected a best fit model based on AIC from the full set of possible combinations of predictor variables using the 'dredge' function from the *MuMIn* R package (10) to avoid overfitting. The best fit model included the following variables: $SA_t * SoCal_t$, $CA_t * SoCal_t$, $E_t$, $P_t$, and the population offset, $log(TotPop_t)$. Lastly, to obtain monthly estimates, we multiplied the total estimate of travel-associated cases for a given tract ($C_t$) by the proportion of cases reported in a given month across the state. Coefficient estimates and p-values for the full and final model predictors are shown in Supplementary Table S3.

To assess performance of our model in predicting the number of dengue cases in other counties across the state, we aggregate our census tract-level predictions to the county level and model reported county-level cases as a function of our county-level predictions using a simple univariate linear model. Reported case counts at the county level between 2010-2023 were obtained from the Centers for Disease Control and Prevention (11). In this dataset, counties reporting between 1 and 4 total cases are assigned a categorical value of '1 to 4'. For our modeling purposes, we conservatively assume that these counties each reported only one case. We find strong agreement between predicted and reported cases at the county level, and good model fit ($R^2$ = 0.94; Supplementary Figure S2).

Although we estimate vector presence and temperature-dependent suitability for transmission at future time points, we only estimate travel-associated cases at present given a lack of comprehensive, fine-scaled projections on socioeconomic and demographic characteristics for California.

3. *Risk metrics*

Our risk metrics combined monthly estimates of vector presence, temperature-dependent transmission suitability, and (for eco-epidemiological risk) travel-associated cases through raster multiplication of these components. In doing so, each component is inherently weighted equally, as has been done in prior multiplicative models (12), but which may not necessarily reflect biological realism. We chose to use this approach rather than estimate different weights for each component (eg, obtaining coefficient estimates through regressing observed cases against each component) to avoid overfitting given the limited number of observed cases at the time of analysis (ie, 8 unique instances).

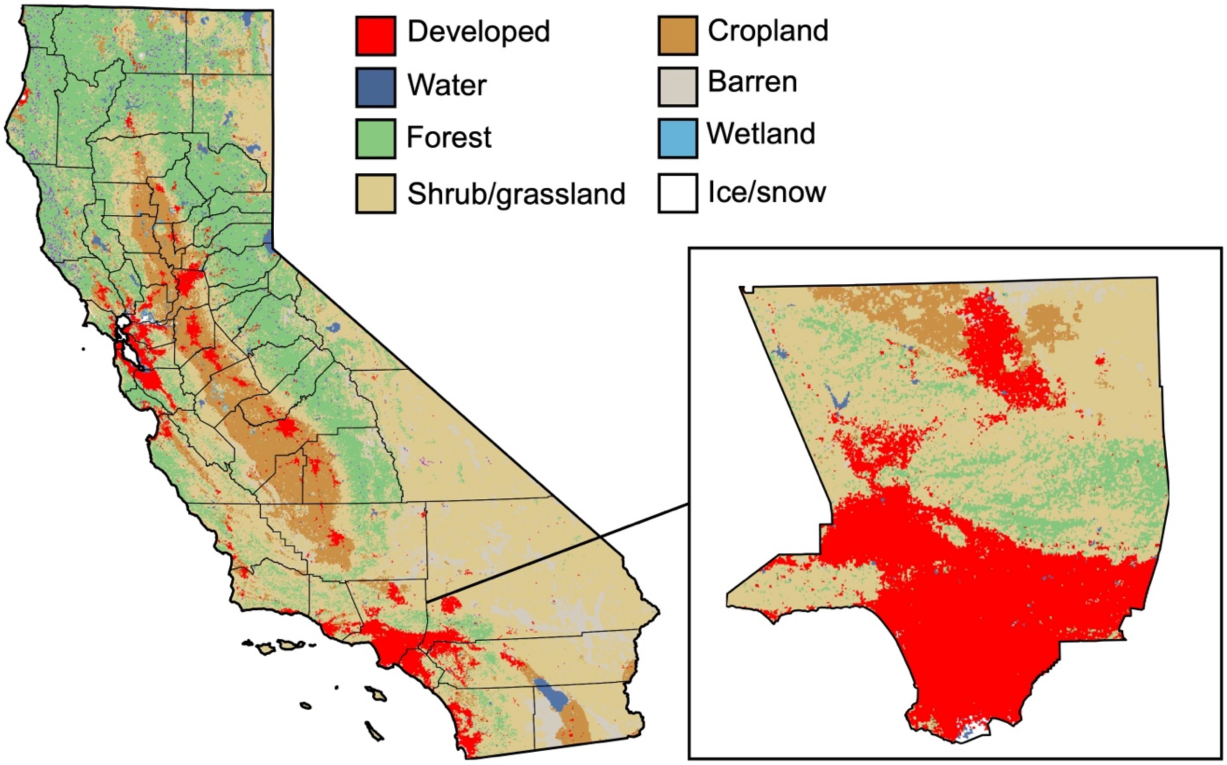


**Supplementary Figure S1.** Land cover designations for California (left) and Los Angeles County specifically (inset). Data were obtained from the National Land Cover Database and available at an annual, 250 m resolution (plots show data from 2000).

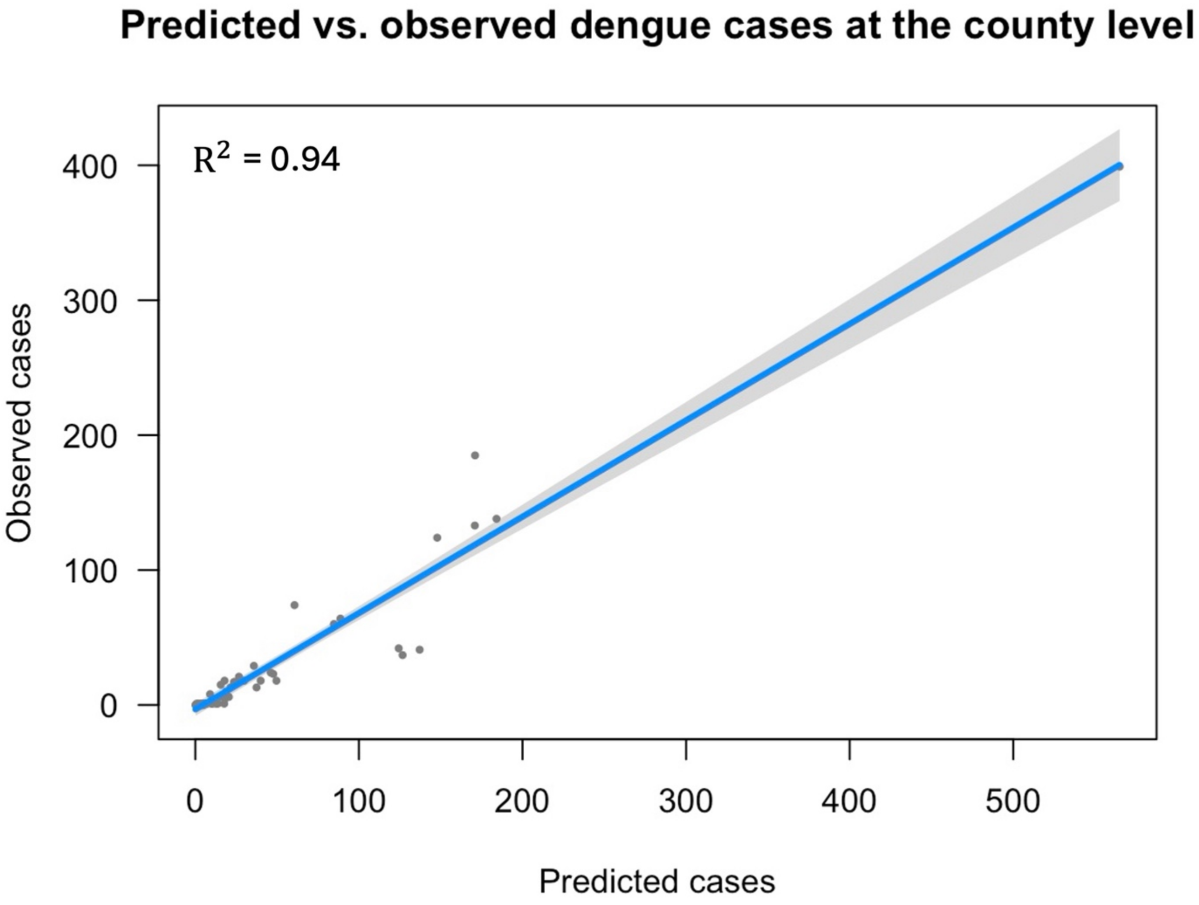


**Supplementary Figure S2.** Comparison of our census tract-level case estimates aggregated by county to the actual number of reported cases per county ($R^2 = 0.94$). Here, cases denote the total number across all months and years (2010-2023) for a given county.

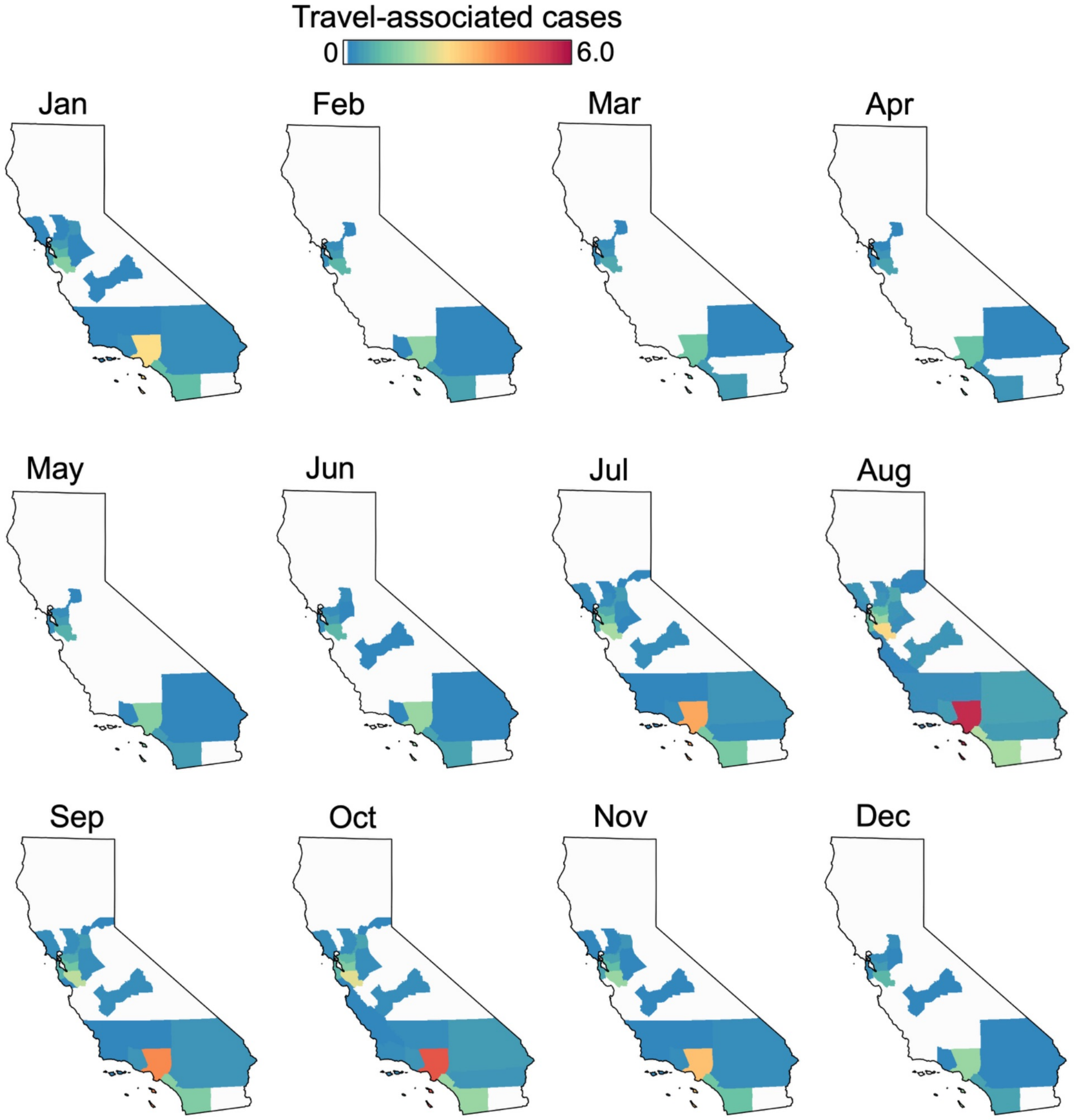


**Supplementary Figure S3.** Monthly counts of travel-associated cases estimated at the county-level (rather than the census tract level). Counts range from 0 to 6.0—the lowest and highest values estimated across all counties and months, respectively.

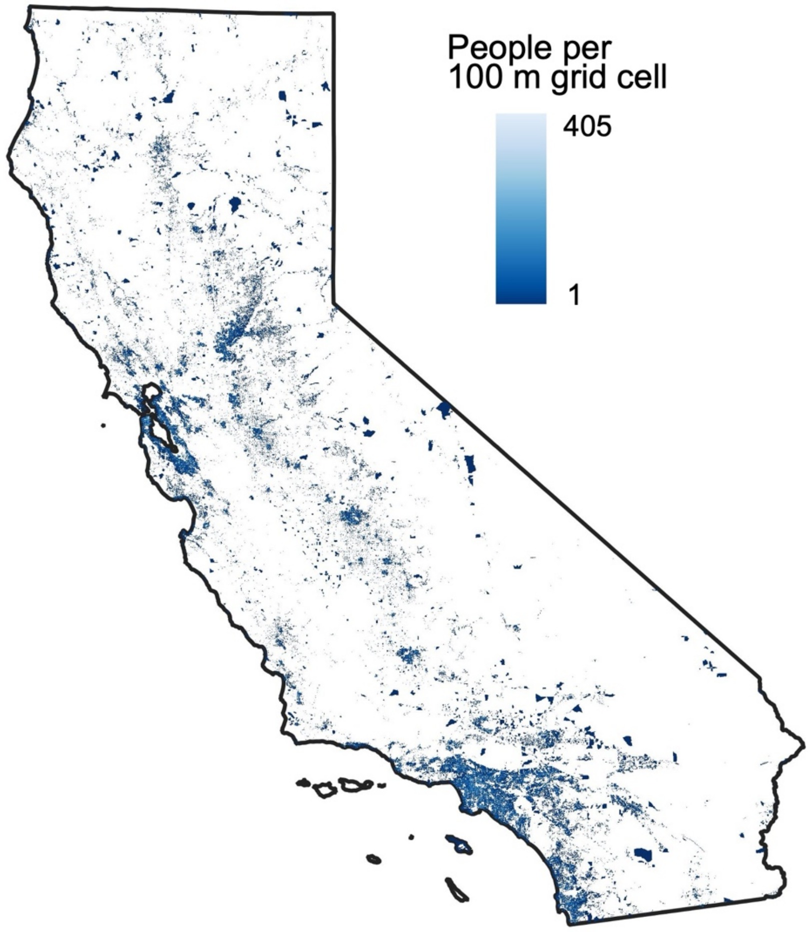


**Supplementary Figure S4.** California population density estimates used in our analysis (13). Pixels denote the number of people per 100 m grid cell. White pixels denote unpopulated areas.

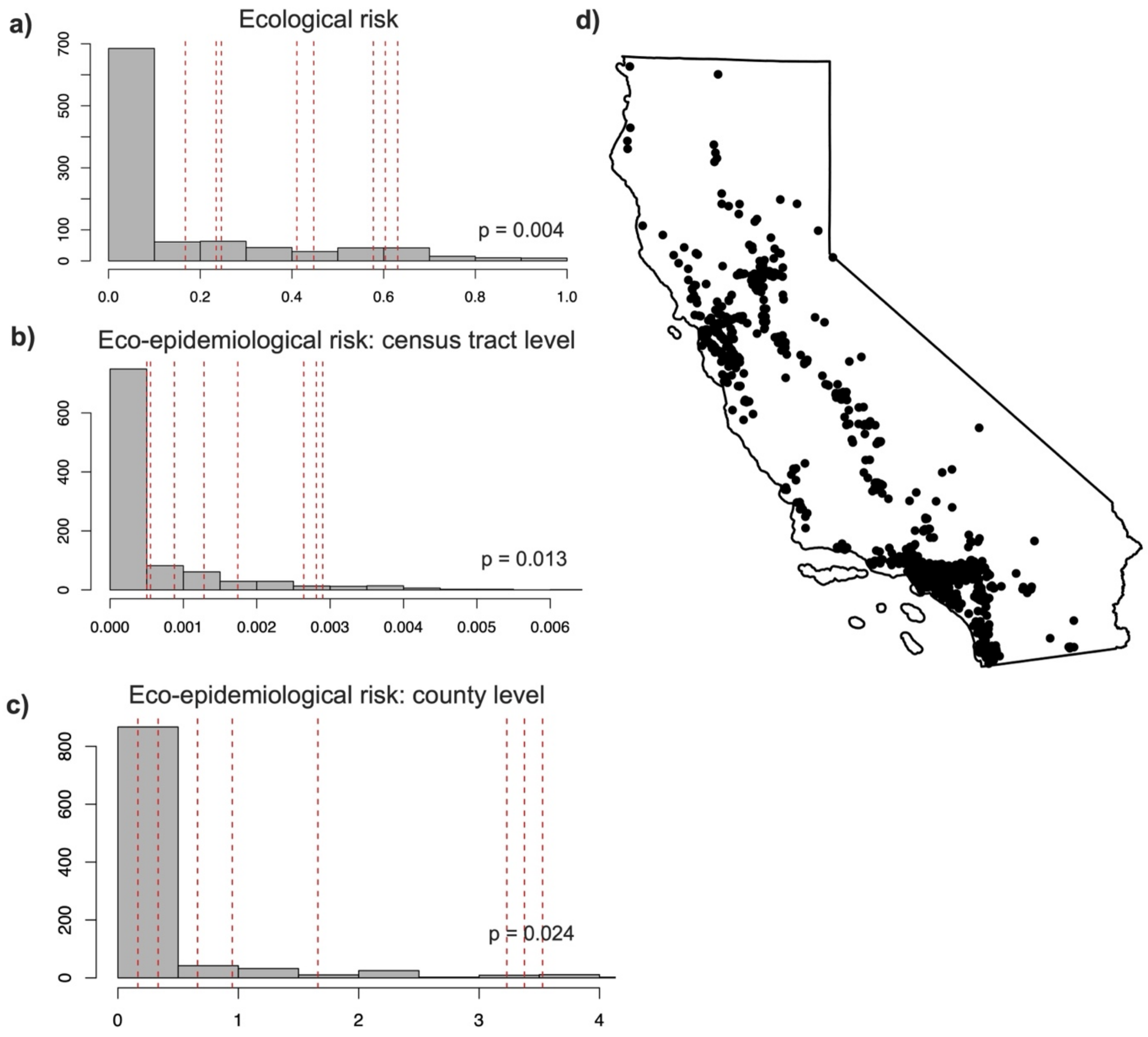


**Supplementary Figure S5.** a-c) Comparison of ecological or eco-epidemiological risk values estimated during observed local transmission (dashed red lines) to estimates from similar months and locations (gray bars see *Risk metric: Definition, calibration, and validation)*. Eco-epidemiological risk estimates incorporated travel-associated cases estimated at either the b) census tract or c) county level. d) Example of population density-weighted random sampling to select 'matched control' locations for comparison with regions where local transmission occurred.

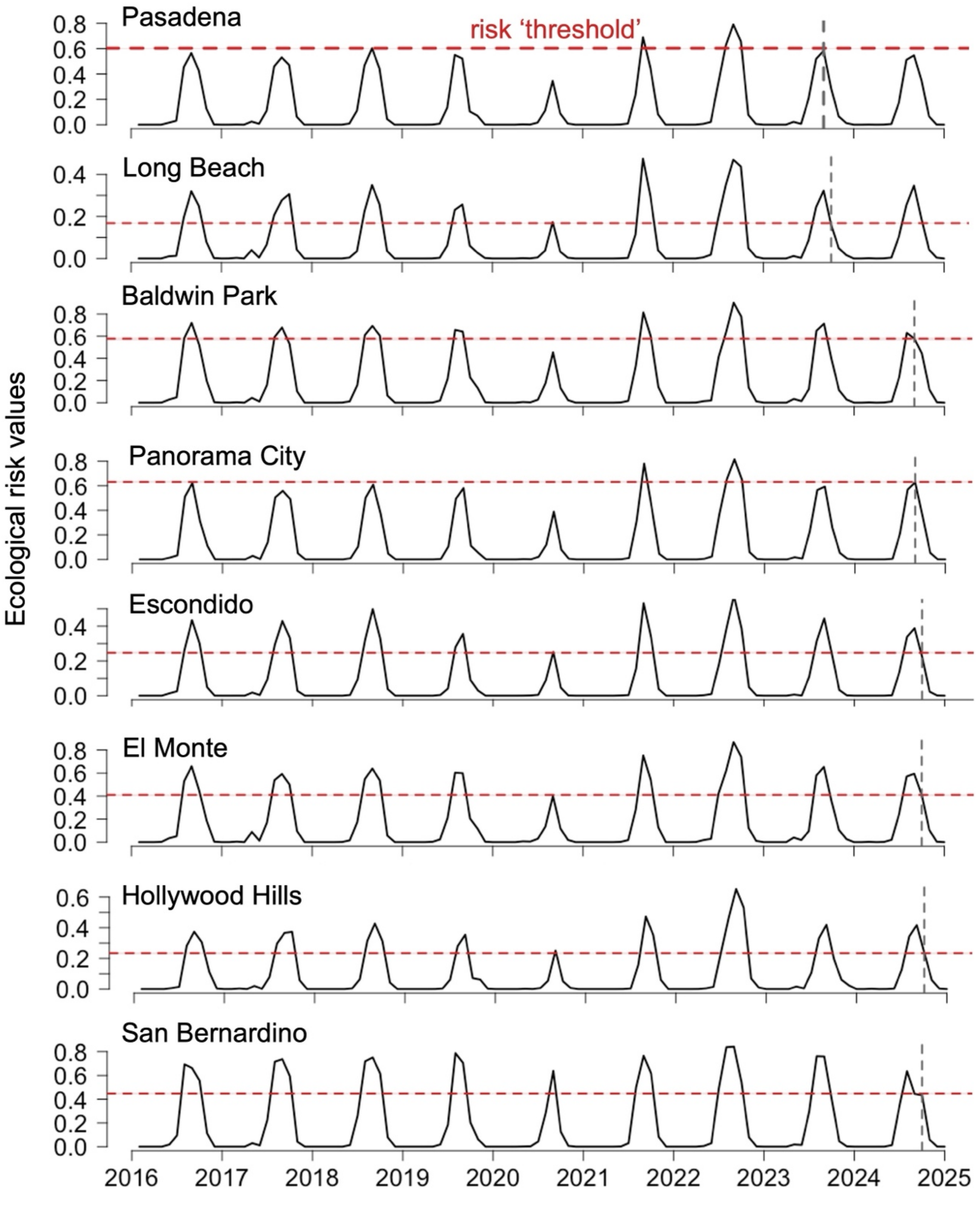


**Supplementary Figure S6.** Monthly values of ecological risk (black) for each of the eight locations in which local dengue transmission was reported in California in 2023-2024. The gray, vertical dashed line in each plot denotes the month in which local transmission putatively occurred (1-2 months prior to case reporting). The red, horizontal dashed line and shading in each plot denote the mean and 95% confidence interval, respectively, of ecological risk during that month. Note that *Ae. aegypti* was first detected in California (Fresno, Madera, and San Mateo counties) in 2013.

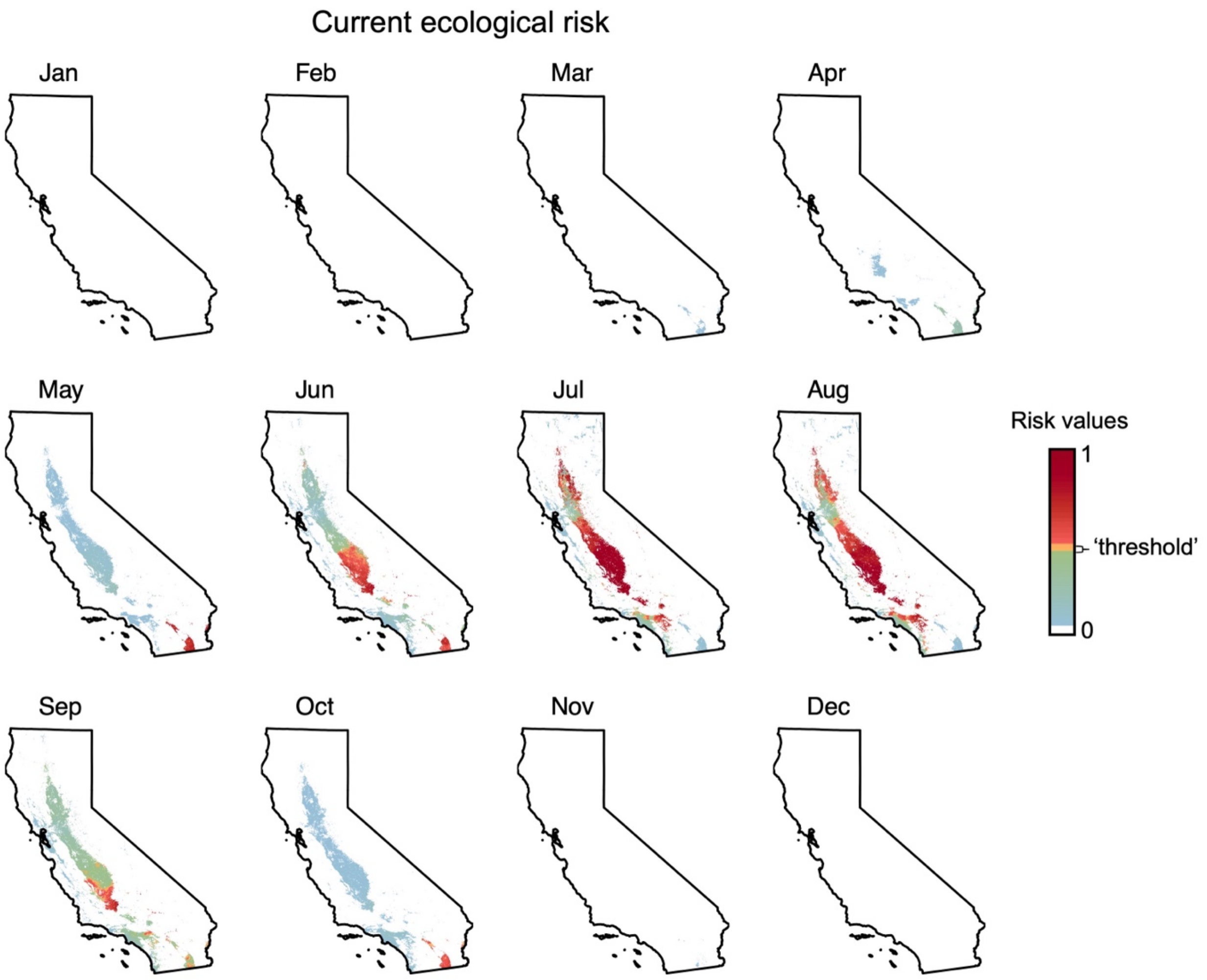


**Supplementary Figure S7.** Monthly estimates of current (2010-2020) ecological risk across California. On the color scale, white denotes zero risk, blue-green denotes low risk (non-zero but below the 'threshold'), yellow-orange denotes risk values estimated during observed local transmission in 2023-2024 ('threshold'), and red denotes risk above this threshold. Note, the scale extends to 1 but the highest estimate across all months was 0.96.

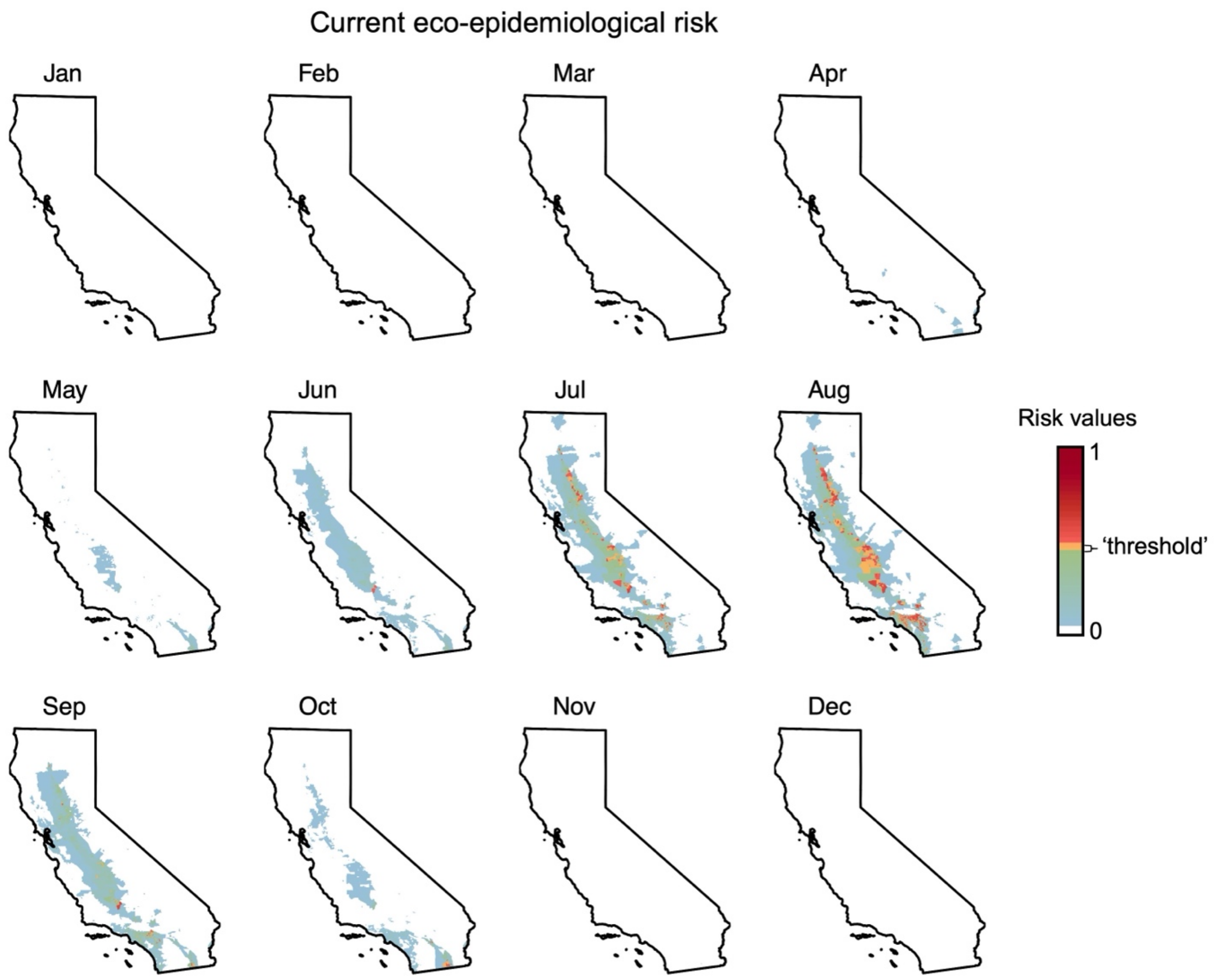


**Supplementary Figure S8.** Monthly estimates of current (2010-2020) eco-epidemiological risk across California. On the color scale, white denotes zero risk, blue-green denotes low risk (non-zero but below the 'threshold'), yellow-orange denotes risk values estimated during observed local transmission in 2023-2024 ('threshold'), and red denotes risk above this threshold.

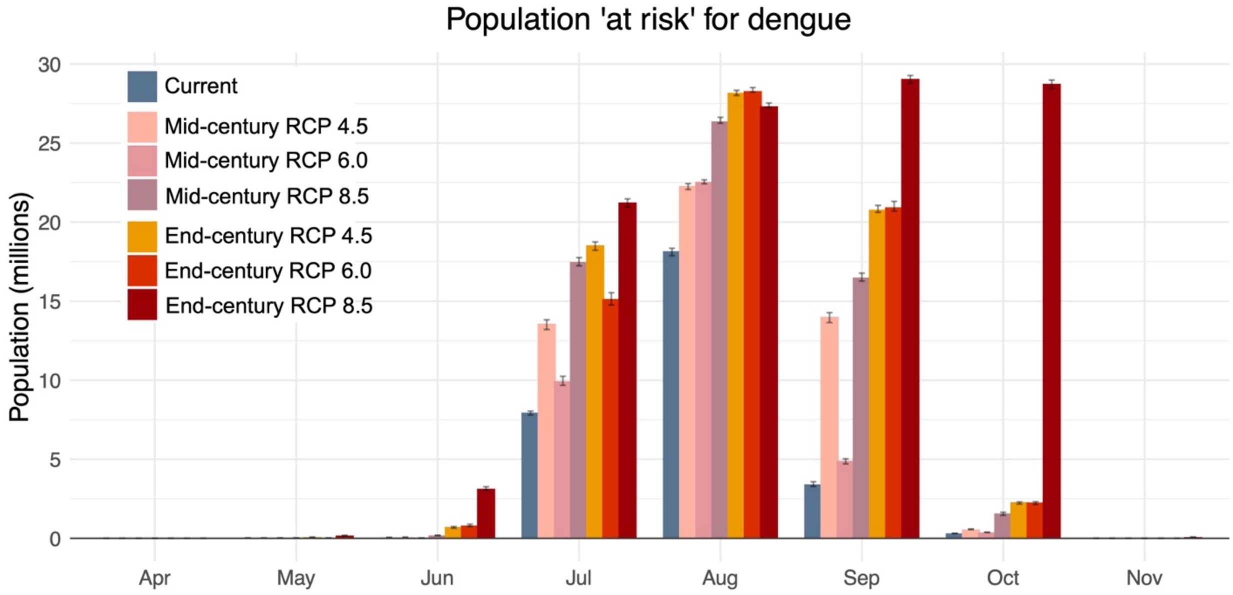


**Supplementary Figure S9.** Population at eco-epidemiological risk for dengue transmission under varying climate warming scenarios. Bars denote the number of California residents (in millions) living within a census tract where the mean eco-epidemiological risk for dengue transmission is at or above the 'threshold' based on observed local transmission. The threshold is based on the mean risk value estimated for census tracts and months with reported local transmission, while error bars denote estimates based on the lower and upper bounds of the 95% confidence interval around that mean. Estimates are shown for the current period (2010-2020) and under two moderate and one upper climate warming scenario (RCP 4.5, 6.0, and 8.5, respectively) for mid-century (2040-2050) and end-of-century (2090-2100).

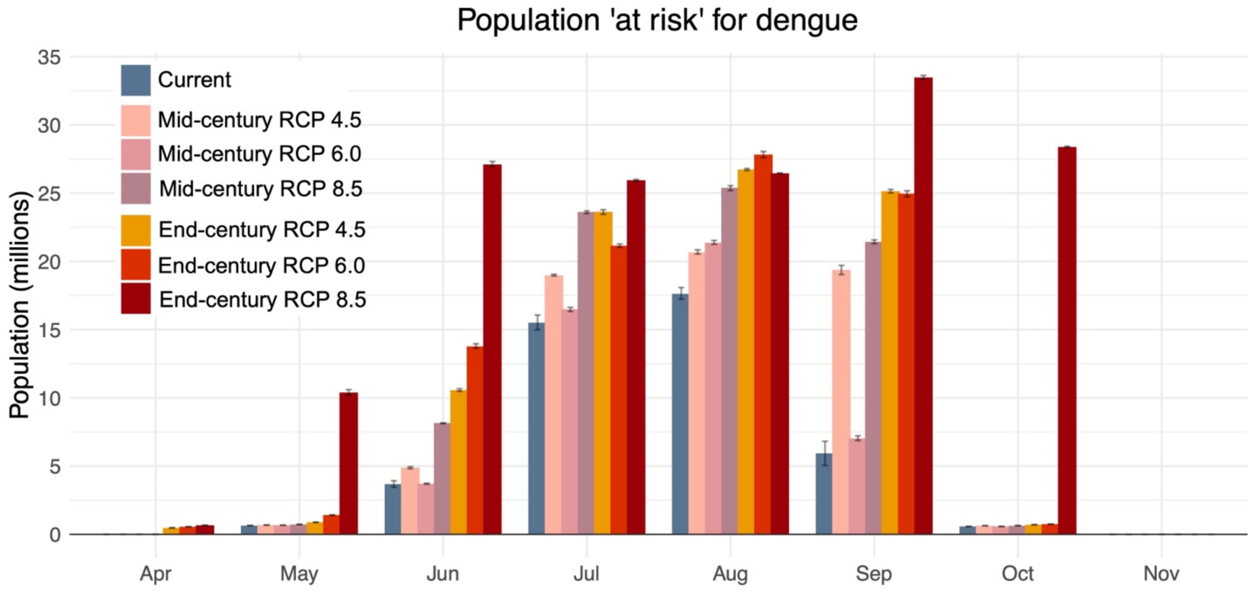


**Supplementary Figure S10.** Population at ecological risk for dengue transmission under varying climate warming scenarios. Bars denote the number of California residents (in millions) living within 1 km of a region where the eco-epidemiological risk for dengue transmission is at or above the 'threshold' based on observed local transmission. The threshold is based on the mean risk value estimated for census tracts and months with reported local transmission, while error bars denote estimates based on the lower and upper bounds of the 95% confidence interval around that mean. Estimates are shown for the current period (2010-2020) and under two moderate and one upper climate warming scenario (RCP 4.5, 6.0, and 8.5, respectively) for mid-century (2040-2050) and end-of-century (2090-2100).

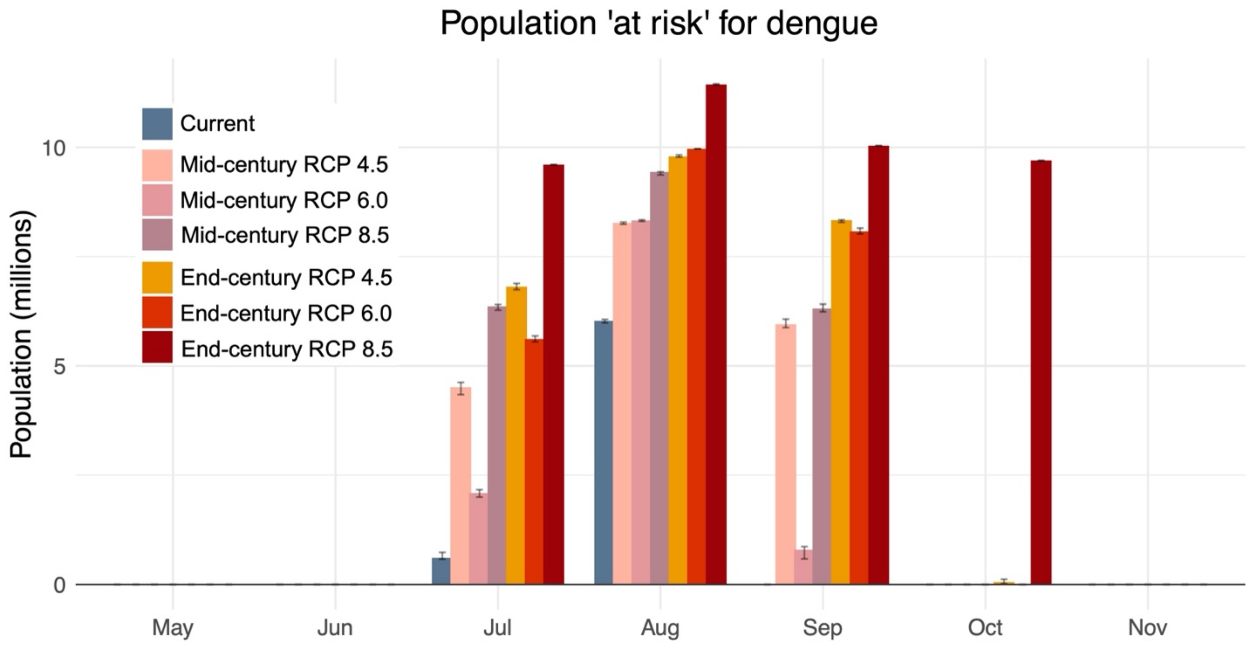


**Supplementary Figure S11.** Population at eco-epidemiological risk for dengue transmission under varying climate warming scenarios based on travel-associated cases estimated at the county level (rather than the census tract level as in the main model specification). Bars denote the number of California residents (in millions) living within a census tract where the mean eco-epidemiological risk for dengue transmission is at or above the 'threshold' based on observed local transmission. The threshold is based on the mean risk value estimated for census tracts and months with reported local transmission, while error bars denote estimates based on the lower and upper bounds of the 95% confidence interval around that mean. Estimates are shown for the current period (2010-2020) and under two moderate and one upper climate warming scenario (RCP 4.5, 6.0, and 8.5, respectively) for mid-century (2040-2050) and end-of-century (2090-2100). We note that the risk estimates are substantially lower than those calculated using census tract-level estimates of travel-associated cases (Figure 3 in the main text), likely because the 'threshold' was now higher as local transmission occurred in three of the five most populous California counties (ie, Los Angeles, San Diego, San Bernardino) (13).

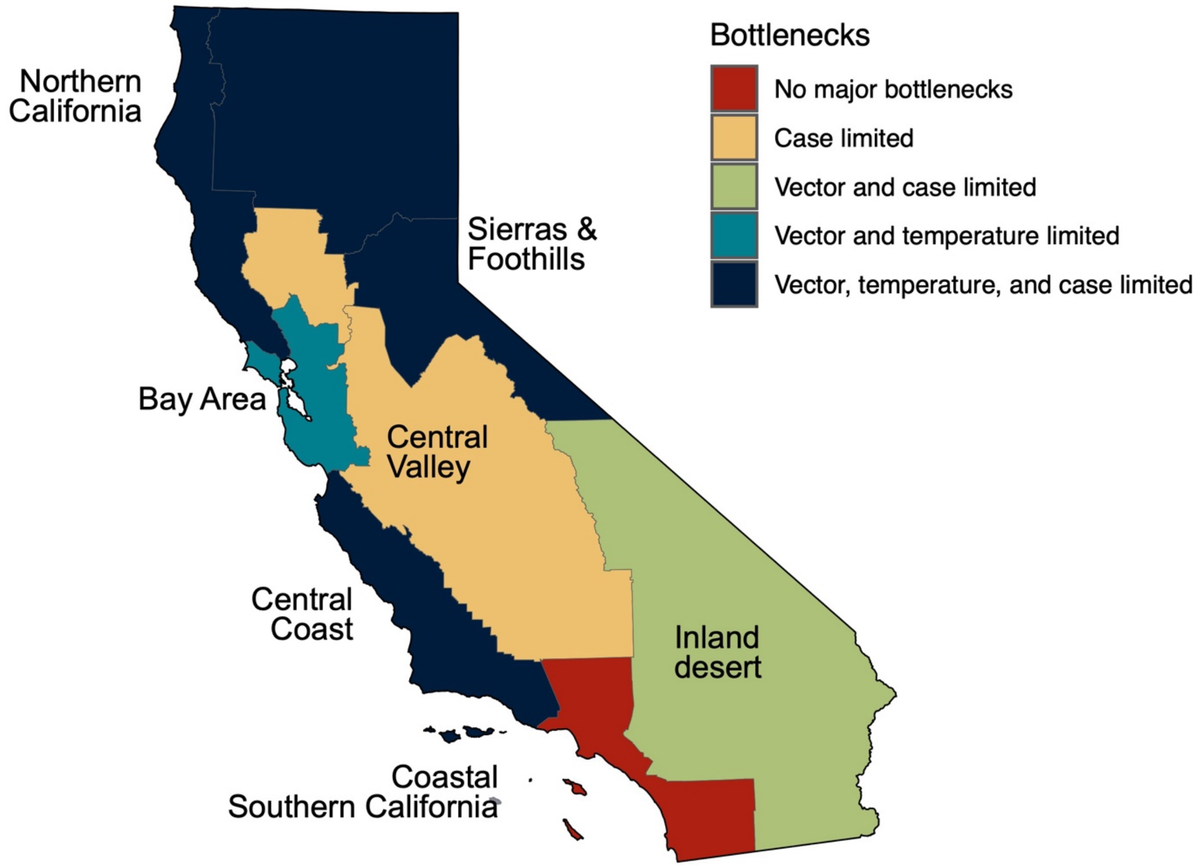


**Supplementary Figure S12.** Regional bottlenecks to transmission under current conditions. Bottlenecks were determined based on regional averages during the peak transmission months (August and September) for each of the components needed for local transmission: vector presence, temperature suitability for pathogen transmission ($R_0(T)$), and viral introductions via travel-associated cases. Regions were considered 'limited' for a component if: vector presence probability < 0.20, $R_0(T) < 0.29$, and/or travel-associated cases < 1.36 cases per month across the region. These classifications are based on the minimum regional-level averages during observed local transmission and are mainly intended to simplify and visualize general patterns rather than capture true biological or epidemiological thresholds. Further, we note that regions with vector presence above the threshold (eg, the Bay Area) may still be vector limited due to extensive vector control efforts.

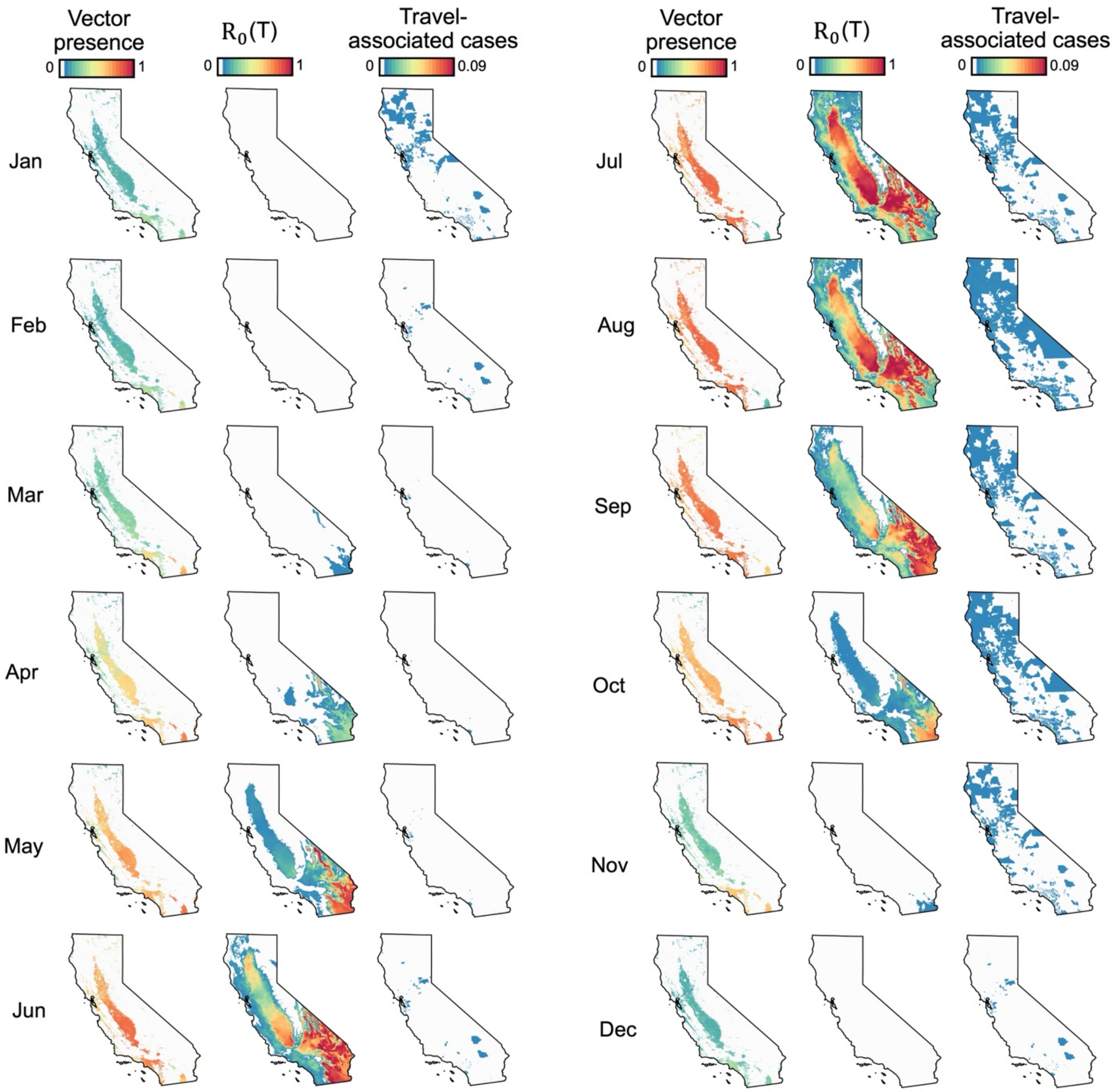


**Supplementary Figure S13.** Monthly values for each of the components included in our model of local dengue transmission risk: vector presence probability (left), temperature-dependent transmission suitability ($R_0(T)$) (center), and travel-associated cases (right). Values denote monthly estimates aggregated across 2010-2020 with a 270 m resolution for vector presence and $R_0(T)$, and census tract-level resolution for travel-associated cases. Specifically, travel-associated cases are expressed as cases per tract-month and range from 0 to 0.09—the lowest and highest values estimated across all census tracts and months, respectively.

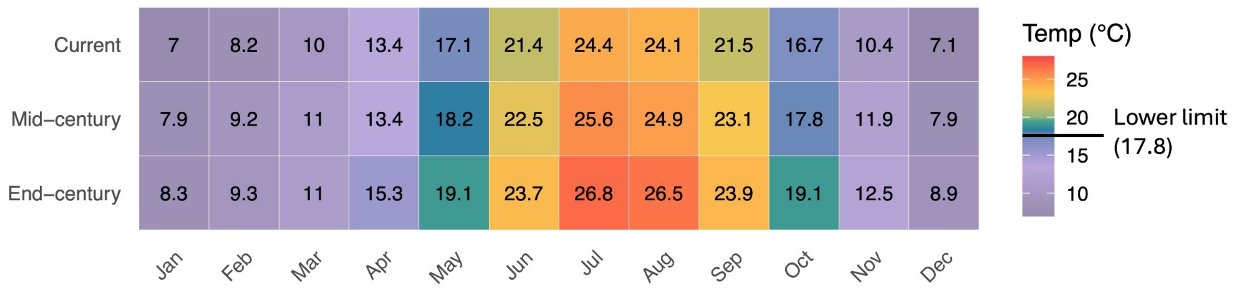


**Supplementary Figure S14.** Monthly temperatures averaged across the state for current (2010-2020), mid-century (2040-2050) and end-of-century (2090-2100) time periods under a moderate climate warming scenario (RCP 4.5). Purple denotes temperatures below the lower thermal limit for DENV transmission by *Aedes aegypti* (17.8°C) as estimated by Mordecai et al. 2017 (14), while the blue-red gradient denotes temperatures above this value.

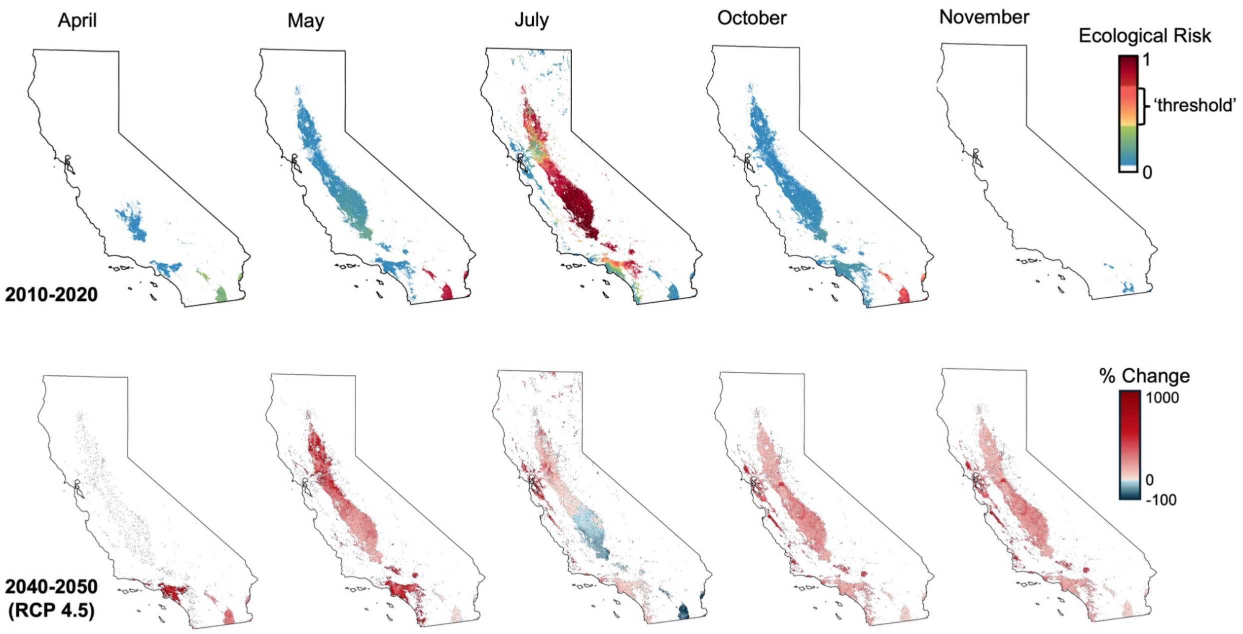


**Supplementary Figure S15.** Current (top panel) and mid-century (bottom panel) ecological risk of transmission. Estimates are shown for four shoulder months (April, May, October, and November) and one peak month (July). Colors in the top panel denote values of our estimated risk metric wherein white denotes zero risk, blue-green denotes low risk (ie, non-zero but below the risk 'threshold'), and yellow-red denotes risk at or above the 'threshold' from observed local transmission. Colors in the bottom panel denote the percent change from current risk, calculated on a per-pixel (270 m) basis.

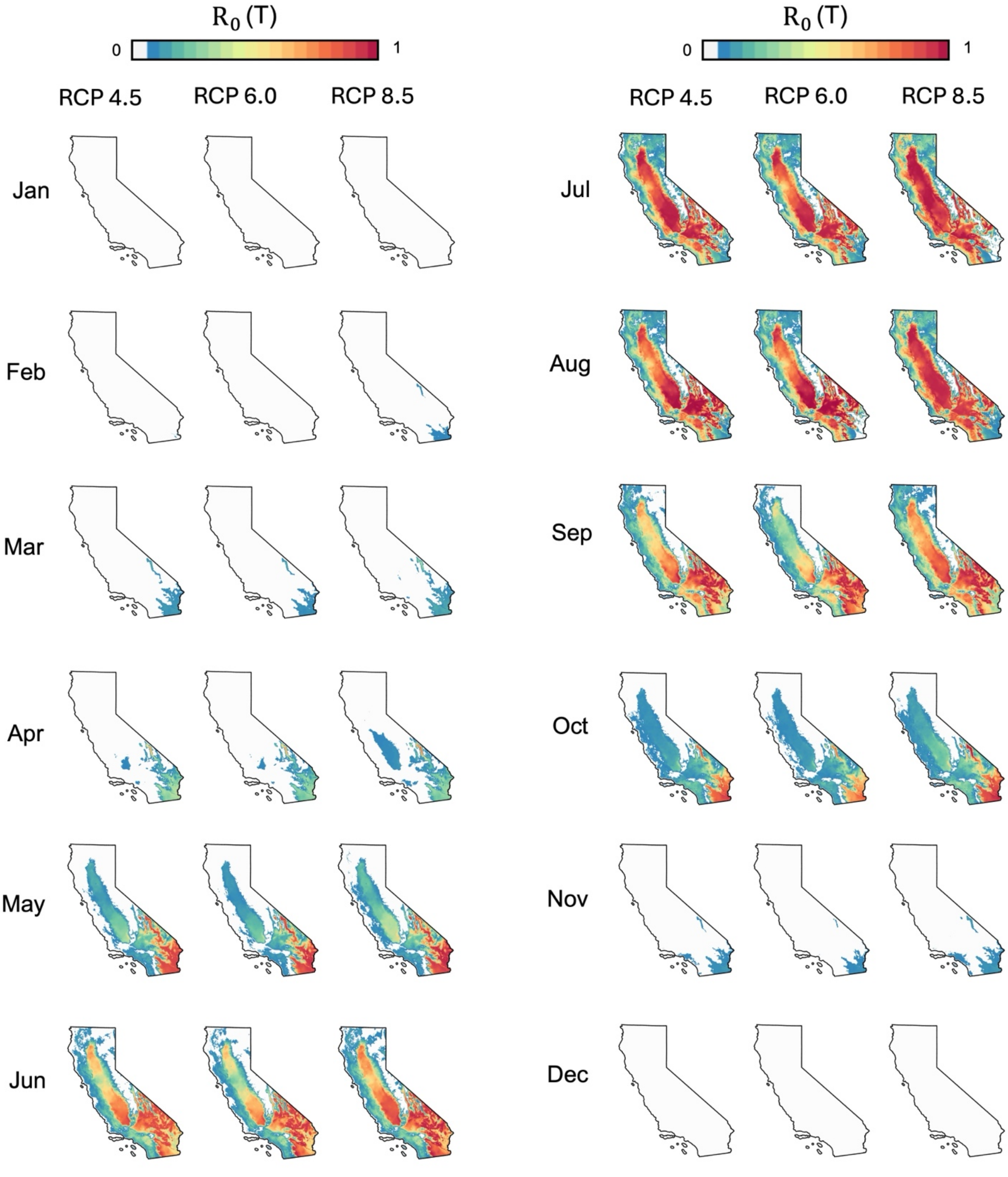


**Supplementary Figure S16.** Monthly estimates of temperature-dependent transmission suitability ($R_0$(T)) for mid-century (2040-2050) under three warming scenarios: RCP 4.5 (left), 6.0 (center), and 8.5 (right).

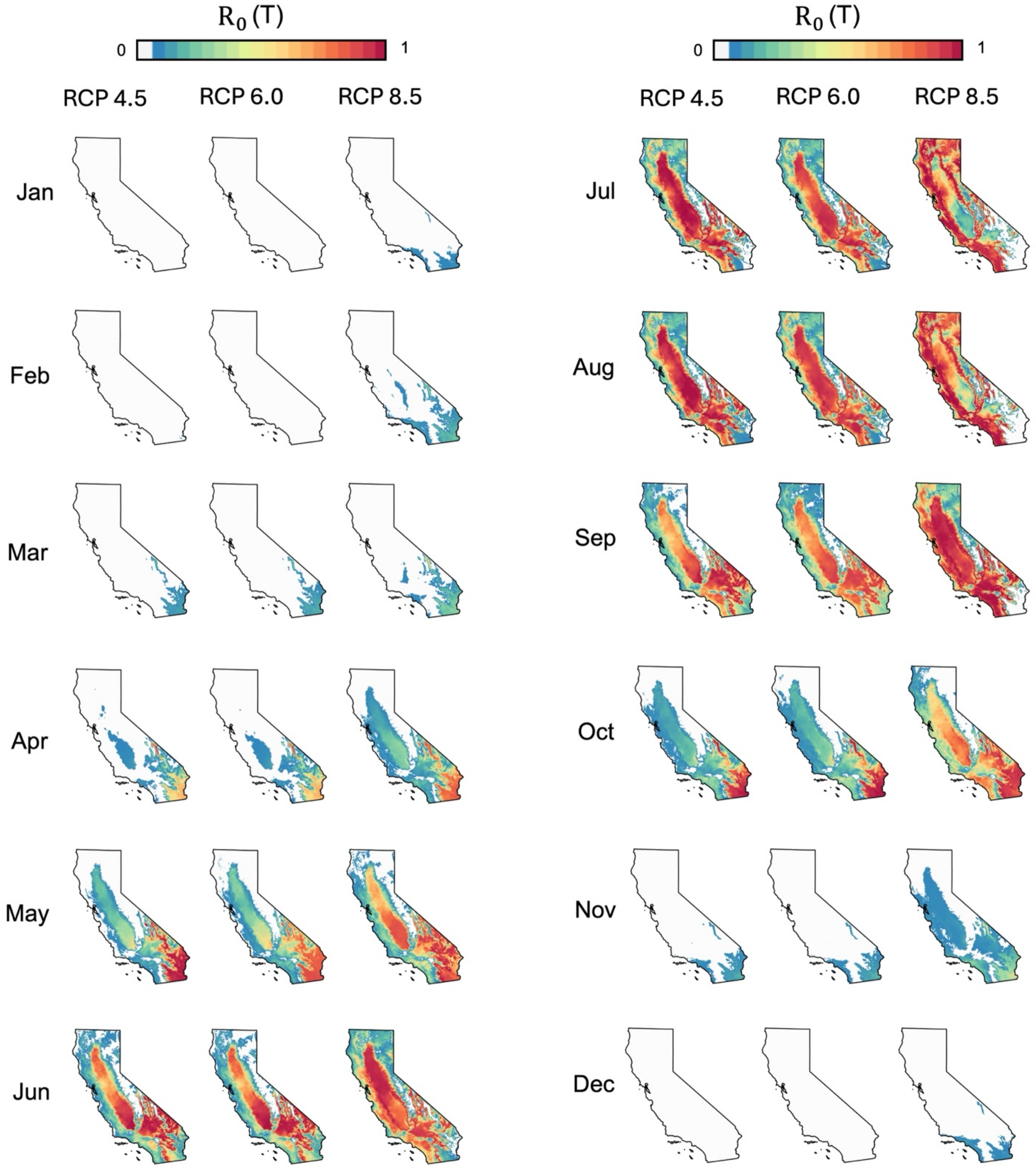


**Supplementary Figure S17.** Monthly estimates of temperature-dependent transmission suitability ($R_0(T)$) for end-of-century (2090-2100) under three warming scenarios: RCP 4.5 (left), 6.0 (center), and 8.5 (right).

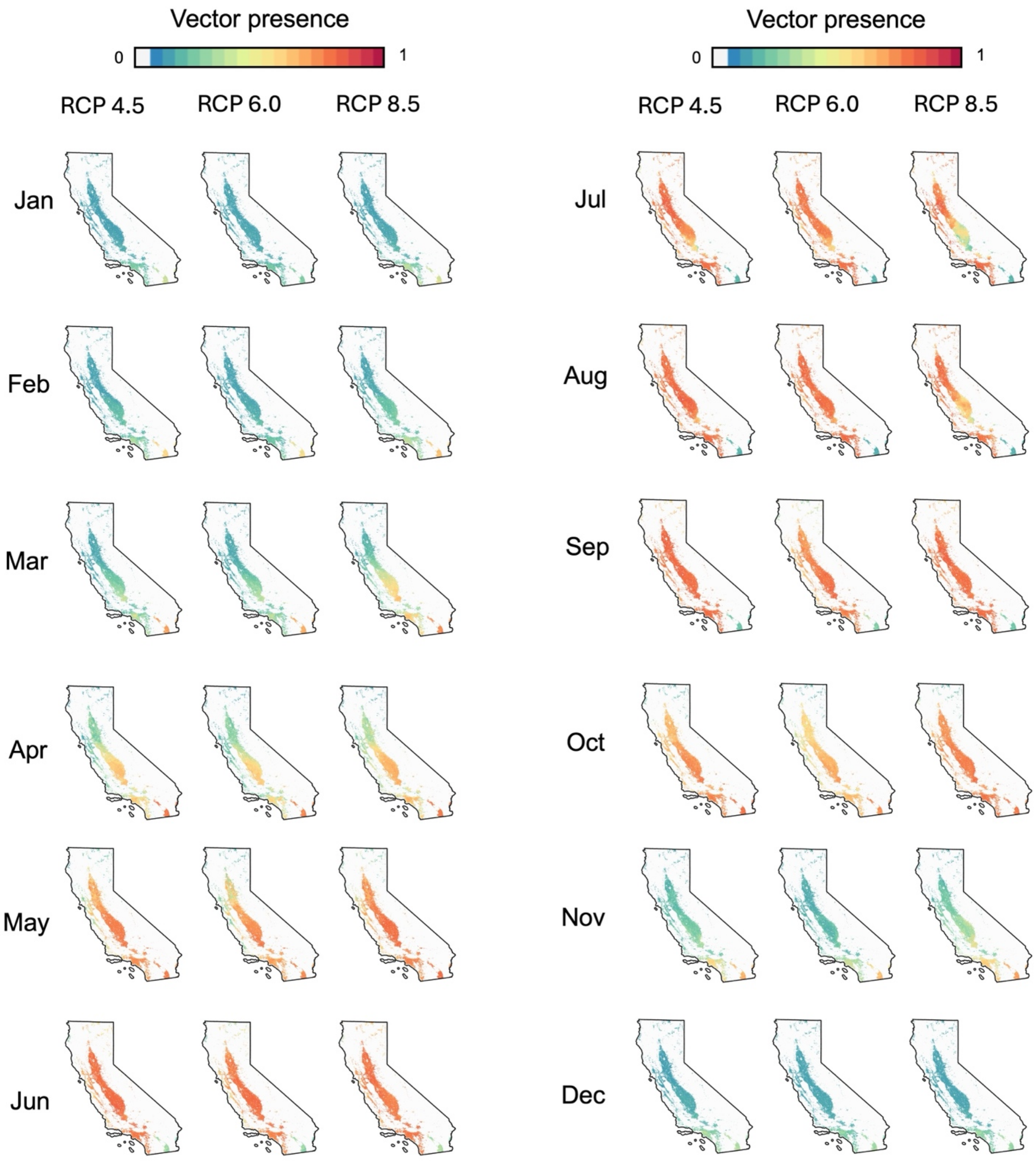


**Supplementary Figure S18.** Monthly estimates of vector presence (expressed as a probability) for mid-century (2040-2050) under three warming scenarios: RCP 4.5 (left), 6.0 (center), and 8.5 (right).

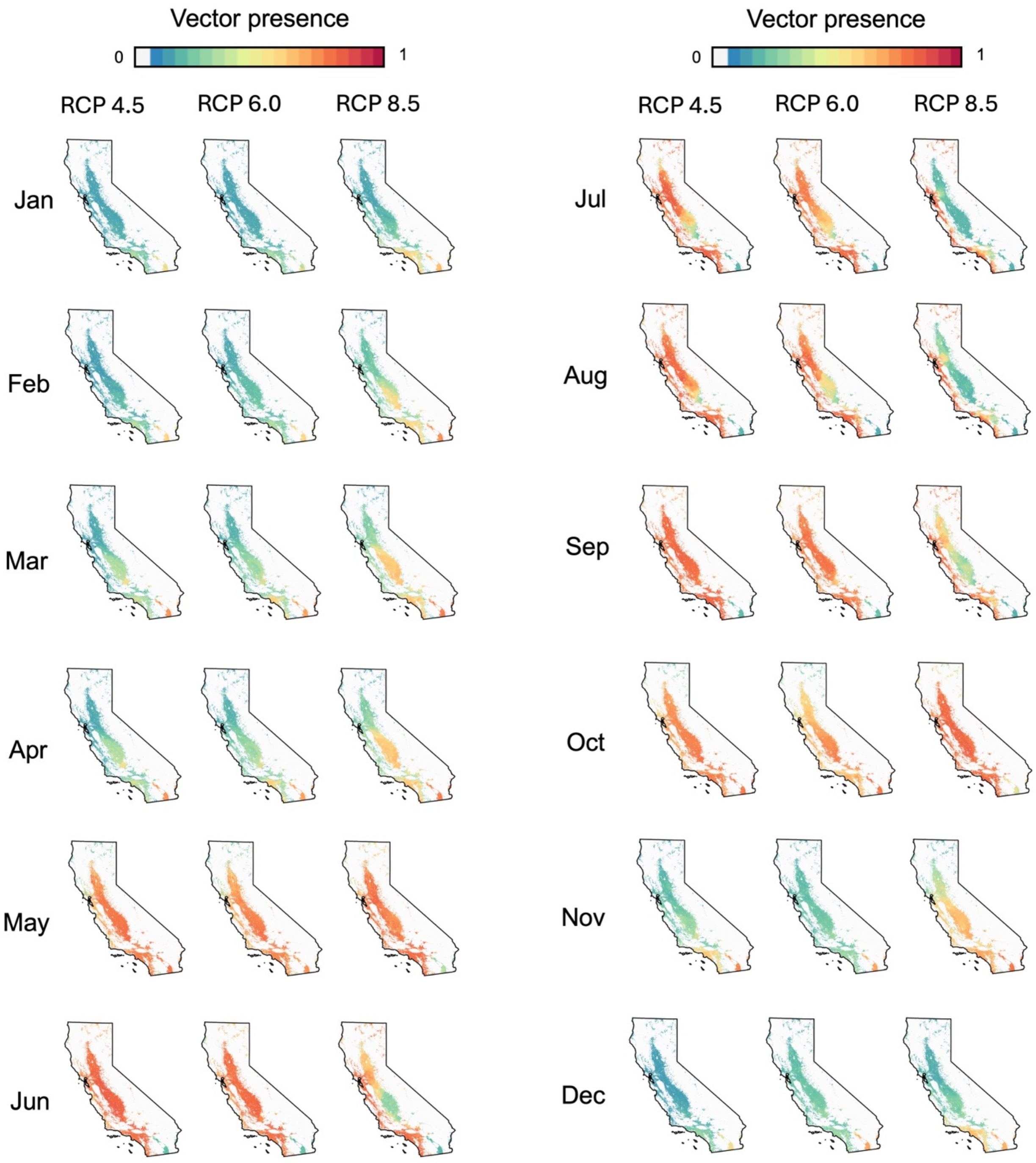


**Supplementary Figure S19.** Monthly estimates of vector presence (expressed as a probability) for end-of-century (2090-2100) under three warming scenarios: RCP 4.5 (left), 6.0 (center), and 8.5 (right).

**Supplementary Table S1.** Mapping of race/ethnicity to world regions and percent of California cases originating from exposure in that region. Population counts by racial and ethnic groups were obtained from the U.S. Census Bureau Detailed Demographic and Housing Characteristics File A (DHC-A) and were mapped to 14 world regions, according to the United Nations geographic regions (UN Statistics Division, 2024). Information on exposure by region was obtained from case surveillance by the California Department of Public Health.

| World Region | Race/Ethnicity Group Label | % Cases |
|---|---|---|
| Polynesia | Polynesian (alone or in any combination), Samoan (alone or in any combination), Tongan (alone or in any combination), Walllisian and Futunan (alone or in any combination), Tongan (alone or in any combination), Tokelauan (alone or in any combination), French Polynesian (alone or in any combination), Cook Islander alone, Tuvaluan alone, Easter Islander alone, Niuean alone | 0.47% |
| North Africa | Sudanese (alone or in any combination), Middle Eastern or North African (alone or in any combination), Algerian (alone or in any combination), Moroccan (alone or in any combination), Egyptian (alone or in any combination), Libyan (alone or in any combination) | 0.17% |
| East Africa | Eritrean (alone or in any combination), Ethiopian (alone or in any combination), Somali (alone or in any combination), Ugandan (alone or in any combination), Kenyan (alone or in any combination), South Sudanese (alone or in any combination), Zambian alone, Djiboutian alone, Malawian alone, Tanzanian (alone or in any combination), Somali alone, Mozambican (alone or in any combination), Rwandan (alone or in any combination), Zimbabwean (alone or in any combination), Burundian (alone or in any combination) | 0.37% |
| West Africa | Nigerian (alone or in any combination), Ghanian (alone or in any combination), Ivoirian alone, Liberian (alone or in any combination), Malian (alone or in any combination), Togolese alone, Sierra Leonean (alone or in any combination), Burkinabe alone, Guinean (alone or in any combination), Bisseau-Guinean (alone or in any combination), Senegalese (alone or in any combination), Gambian (alone or in any combination), Mauritanian (alone or in any combination), Cabo Verdean (alone or in any combination) | 0.50% |
| Middle Africa | Cameroonian (alone or in any combination), Congolese (alone or in any combination), Angolan (alone or in any combination), Equatorial Guinean alone, Chadian alone, Gabonese alone | 0.03% |
| Southern Africa | South African (alone or in any combination), Motswana (alone or in any combination), Swazi (alone or in any combination), Namibian alone | 0.15% |
| Caribbean | Puerto Rican, Cuban, Caribbean Hispanic, Dominican, Other Caribbean Hispanic, Jamaican (alone or in any combination), Antiguan and Barbudan alone, Haitian (alone or in any combination), Caribbean (alone or in any combination), Trinidadian and Tobagonian (alone or in any combination), Montserratian (alone or in any combination), Barbadian (alone or in any combination), Kittian and Nevisian (alone or in any combination), British Virgin Islands alone, Bahamian (alone or in any combination), U.S. Virgin Islander alone, Vincentian alone, West Indian (alone or in any combination), Anguillan alone | 4.00% |

| | | |
|---|---|---|
| Central America | Mexican, Salvadoran, Nicaraguan, Guatemalan, Central American, Honduran, Other Central American, Panamanian, Costa Rican, Belizean (alone or in any combination) | 49.00% |
| South America | South American, Colombian, Argentinean, Peruvian, Ecuadorian, Venezuelan, Chilean, Bolivian, Other South American, Uruguayan, Paraguayan, Brazilian (alone or in any combination), Guyanese (alone or in any combination) | 3.14% |
| Melanesia | Fijian (alone or in any combination), Melanesian (alone or in any combination), Rotuman (alone or in any combination), Nauruan (alone or in any combination), Soloman Islander alone, Papua New Guinean (alone or in any combination), New Caledonian (alone or in any combination) | 0.22% |
| Western Asia | Iranian (alone or in any combination), Israeli (alone or in any combination), Lebanese (alone or in any combination), Yemeni (alone or in any combination), Arab (alone or in any combination), Armenian (alone or in any combination), Turkish (alone or in any combination), Iraqi (alone or in any combination), Palestinian (alone or in any combination), Jordanian (alone or in any combination), Syrian (alone or in any combination), Saudi (alone or in any combination), Omani alone, Kurdish (alone or in any combination), Qatari (alone or in any combination), Assyrian (alone or in any combination), Georgian (alone or in any combination), Kuwaiti (alone or in any combination), Chaldean (alone or in any combination), Cypriot (alone or in any combination), Syriac alone, Azerbaijani (alone or in any combination) | 0.17% |
| East Asia | East Asian, Chinese except Taiwanese (alone or in any combination), Japanese (alone or in any combination), Korean (alone or in any combination), East Asian (alone or in any combination), Taiwanese (alone or in any combination), Mongolian (alone or in any combination) | 0.25% |
| South Asia | Asian Indian (alone or in any combination), South Asian (alone or in any combination), Pakistani (alone or in any combination), Afghan (alone or in any combination), Nepalese (alone or in any combination), Sri Lankan (alone or in any combination), Bhutanese (alone or in any combination), Maldivian alone, Bangladeshi (alone or in any combination), Sikh (alone or in any combination), Kuki alone, Sindhi (alone or in any combination), Mizo (alone or in any combination), Pashtun (alone or in any combination) | 25.39% |
| Southeast Asia | Filipino (alone or in any combination), Vietnamese (alone or in any combination), Southeast Asian (alone or in any combination), Indonesian (alone or in any combination), Cambodian (alone or in any combination), Burmese (alone or in any combination), Laotian (alone or in any combination), Thai (alone or in any combination), Burmese (alone or in any combination), Mien (alone or in any combination), Hmong (alone or in any combination), Malaysian (alone or in any combination), Lahu alone, Tai Dam (alone or in any combination), Timorese (alone or in any combination), Bruneian alone, Singaporean (alone or in any combination) | 8.08% |

**Supplementary Table S2.** Estimates of vector presence (expressed as a probability), temperature-dependent transmission suitability ($R_0$(T) scaled from 0 to 1), the number of travel-associated cases per tract-month, ecological risk, and eco-epidemiological risk for each instance of local transmission reported in California in 2023-2024. The month of estimated transmission for a given case was determined based on personal communication with city and state public health officials (M. Feaster, City of Pasadena Public Health Department, personal communication, October 9, 2024).

| Location | Month reported | Month of estimated transmission | $R_0$(T) | Vector presence | Travel-associated cases | Eco. risk | Eco-epi risk |
|---|---|---|---|---|---|---|---|
| Pasadena | Oct 2023 | Aug 2023 | 0.626 | 0.964 | 0.0044 | 0.604 | 0.0026 |
| Long Beach | Oct 2023 | Sep 2023 | 0.156 | 0.865 | 0.0032 | 0.168 | 0.0005 |
| Baldwin Park | Sept 2024 | Aug 2024 | 0.727 | 0.816 | 0.0048 | 0.578 | 0.0027 |
| Panorama City | Sept 2024 | Aug 2024 | 0.680 | 0.910 | 0.0046 | 0.630 | 0.0028 |
| Escondido | Oct 2024 | Sep 2024 | 0.341 | 0.766 | 0.0035 | 0.246 | 0.0009 |
| El Monte | Oct 2024 | Sep 2024 | 0.457 | 0.851 | 0.0031 | 0.411 | 0.0012 |
| Hollywood Hills | Oct 2024 | Sep 2024 | 0.332 | 0.652 | 0.0022 | 0.235 | 0.0005 |
| City of San Bernardino | Nov 2024 | Sep 2024 | 0.187 | 0.952 | 0.0040 | 0.447 | 0.0018 |

**Supplementary Table S3.** Summary statistics from the Poisson generalized linear model estimating census tract-level travel-associated cases. Incidence Rate Ratios correspond to the exponentiated coefficient estimates and represent the relative change in the predicted number of cases given a one-unit increase in the predictor. The left- and right-hand side of the table denote the full and final model, respectively.

| | | | dengue cases | | | |
|---|---|---|---|---|---|---|
| *Predictors* | *Incidence Rate Ratios* | *CI* | *p* | *Incidence Rate Ratios* | *CI* | *p* |
| Intercept | 0.00 | 0.00 – 0.00 | **<0.001** | 0.00 | 0.00 – 0.00 | **<0.001** |
| South Asia | 1.06 | 1.01 – 1.11 | **0.024** | 1.06 | 1.02 – 1.09 | **0.001** |
| SoCal | 0.92 | 0.69 – 1.27 | 0.598 | 0.89 | 0.69 – 1.18 | 0.396 |
| SE Asia | 1.02 | 0.91 – 1.14 | 0.706 | | | |
| Central Am | 0.55 | 0.39 – 0.76 | **0.001** | 0.56 | 0.40 – 0.76 | **<0.001** |
| Total Pop Regions | 1.01 | 0.90 – 1.12 | 0.926 | | | |
| Prop Pop Less College | 0.82 | 0.69 – 0.97 | **0.021** | 0.87 | 0.77 – 0.99 | **0.031** |
| Prop Pop Poverty | 1.05 | 0.95 – 1.16 | 0.317 | 1.09 | 0.99 – 1.19 | 0.081 |
| Prop Pop English Not Well | 1.08 | 0.95 – 1.22 | 0.261 | | | |
| Prop Pop Over 16 Unemp | 1.05 | 0.96 – 1.15 | 0.241 | | | |
| South Asia * SoCal | 0.67 | 0.46 – 0.93 | **0.029** | 0.67 | 0.46 – 0.92 | **0.026** |
| SE Asia * SoCal | 0.93 | 0.79 – 1.09 | 0.371 | | | |
| Central Am * SoCal | 2.08 | 1.51 – 2.95 | **<0.001** | 2.04 | 1.51 – 2.84 | **<0.001** |
| Observations | 2886 | | | 2886 | | |
| $R^2$ Nagelkerke | 0.055 | | | 0.055 | | |

**Supplementary References**


1. Phillips SJ, Anderson RP, Schapire RE. Maximum entropy modeling of species geographic distributions. Ecological Modelling. 2006 Jan 25;190(3):231–59.

2. Muscarella R, Galante PJ, Soley-Guardia M, Boria RA, Kass JM, Uriarte M, et al. ENM eval: An R package for conducting spatially independent evaluations and estimating optimal model complexity for MAXENT ecological niche models. McPherson J, editor. Methods Ecol Evol. 2014 Nov;5(11):1198–205.

3. Strobl C, Malley J, Tutz G. An introduction to recursive partitioning: Rationale, application, and characteristics of classification and regression trees, bagging, and random forests. Psychological Methods. 2009;14(4):323–48.

4. Chen T, Guestrin C. XGBoost: A Scalable Tree Boosting System [Internet]. 2016 [cited 2022 June 1]. Available from: https://dl.acm.org/doi/10.1145/2939672.2939785

5. Kraemer MU, Sinka ME, Duda KA, Mylne AQ, Shearer FM, Barker CM, et al. The global distribution of the arbovirus vectors Aedes aegypti and Ae. albopictus. eLife [Internet]. 2015 June 30 [cited 2020 May 8];4. Available from: https://elifesciences.org/articles/08347

6. Sallam M, Fizer C, Pilant A, Whung PY. Systematic Review: Land Cover, Meteorological, and Socioeconomic Determinants of Aedes Mosquito Habitat for Risk Mapping. IJERPH. 2017 Oct 16;14(10):1230.

7. Benitez EM, Ludueña-Almeida F, Frías-Céspedes M, Almirón WR, Estallo EL. Could land cover influence *Aedes aegypti* mosquito populations? Medical Vet Entomology. 2020 June;34(2):138–44.

8. Metzger ME, Hardstone Yoshimizu M, Padgett KA, Hu R, Kramer VL. Detection and Establishment of Aedes aegypti and Aedes albopictus (Diptera: Culicidae) Mosquitoes in California, 2011–2015. Journal of Medical Entomology. 2017 May 1;54(3):533–43.

9. U.S. Census Bureau. 2020 Census Detailed Demographic and Housing Characteristics File A (Detailed DHC-A). 2020.

10. Bartoń K. MuMIn: Multi-Model Inference [Internet]. New York; 2025. Available from: https://CRAN.R-project.org/package=MuMIn

11. Centers for Disease Control and Prevention. Dengue Historic Data (2010-2014). 2025.

12. Childs ML, Nova N, Colvin J, Mordecai EA. Mosquito and primate ecology predict human risk of yellow fever virus spillover in Brazil. Philosophical Transactions of the Royal Society B: Biological Sciences. 2019;374(20180335).

13. Depsky NJ, Cushing L, Morello-Frosch R. High-resolution gridded estimates of population sociodemographics from the 2020 census in California. Vadrevu KP, editor. PLoS ONE. 2022 July 14;17(7):e0270746.

14. Mordecai EA, Cohen JM, Evans MV, Gudapati P, Johnson LR, Lippi CA, et al. Detecting the impact of temperature on transmission of Zika, dengue, and chikungunya using mechanistic models. PLOS Neglected Tropical Diseases. 2017 Apr 27;11(4):e0005568.
